# Operation of Carbon-Concentrating Mechanisms in Cyanobacteria and Algae requires altered poising of the Calvin-Benson cycle

**DOI:** 10.1101/2022.08.23.504937

**Authors:** Haim Treves, Stefan Lucius, Regina Feil, Mark Stitt, Martin Hagemann, Stéphanie Arrivault

## Abstract

Cyanobacteria and eukaryotic algae make a major contribution to global photosynthetic productivity. To cope with the low availability of CO_2_ in aqueous systems they deploy inorganic carbon-concentrating mechanisms (CCMs). These concentrate CO_2_ in microcompartments that contain Rubisco (carboxysomes in cyanobacteria; pyrenoids in green algae). The rest of the Calvin-Benson cycle (CBC) is located outside these microcompartments. We hypothesized that this physical separation requires modified poising of the CBC. Hence, Rubisco is physically separated from the other CBC enzymes outside these microcompartments. To test the hypothesis that this physical separation requires appropriate poising of the CBC, we profiled CBC metabolites under ambient CO_2_ in the cyanobacterium *Synechocystis* sp. PCC 6803 and three eukaryotic algae (*Chlamydomonas reinhardtii, Chlorella sorokiniana, Chlorella ohadii*). Comparison with recently reported profiles for a large set of terrestrial plants revealed that cyanobacteria and green algae have very distinctive CBC metabolite profiles, with low levels of pentose phosphates and, especially, high levels of ribulose 1,5-bisphosphate and 3-phosphoglycerate. We propose that large pools of the substrate and product of Rubisco are required to generate concentration gradients that drive movement into and out of the microcompartments. These observations raise questions about how CBC regulation was modified during the evolution of algal CCMs and their subsequent loss in terrestrial plants, and highlight that operation of CCMs requires co-evolution of the CBC.

**Highlight:** CBC metabolite profiles in the cyanobacterium Synechocystis and in three eukaryotic green algae at ambient CO2 concentration are very different to those in terrestrial plants, probably reflecting the operation of a carboxysome- or pyrenoid-based carbon concentrating mechanism.

## Introduction

At a global scale, photoautotrophic organisms fix 111-117 × 10^15^ grams of carbon (C) per year, with around half of net primary production being aquatic (Field *et al*., 1998; Behrenfeld *et al*., 2001). There is pressing need to improve our understanding of photosynthetic C fixation, both to improve global C models (Ryu *et al*., 2019) and to underpin breeding and engineering of photosynthesis to increase yield (Long *et al*., 2015; Ort *et al*., 2015). Photosynthetic CO_2_ fixation occurs in the Calvin-Benson cycle (CBC) that evolved together with oxygenic photosynthesis in cyanobacteria and was transferred via endosymbiosis into archaeplastida, i.e. algae and plants (Hohmann-Marriott & Blankenship, 2011).

The CBC consists of three sub-processes; fixation of CO_2_ by Rubisco to form two molecules of 3-phosphoglycerate (3PGA), reduction of 3PGA to triose phosphate (triose-P) using NADPH and ATP from the light reactions, and regeneration of ribulose 1,5-bisphosphate (RuBP) from triose-P (Heldt *et al*., 2005). Fixed C is converted to endproducts including carbohydrates, organic acids and amino acids. In addition, Rubisco catalyses a side reaction with O_2_, leading to formation of 2-phosphoglycolate (2PG) (Lorimer, 1981) that must be salvaged via the photorespiratory pathway in which two 2PG molecules are recycled to one 3PGA molecule with concomitant release of CO_2_ and ammonia (Bauwe, 2018; Busch, 2020). The oxygenase reaction would have been suppressed in the high CO_2_/low O_2_ atmosphere that prevailed 2.7 million years ago when the CBC evolved in photosynthetic bacteria (Rasmussen *et al*., 2008). However, oxygenic photosynthesis led to a massive decrease in the atmospheric CO_2_ and increase in the atmospheric O_2_ concentration. In current atmospheric conditions (∼400 ppm CO_2_, 21% O_2_), about every fourth reaction of Rubisco is with O_2_ (Osmond, 1981; Sharkey, 1988; Hagemann *et al*., 2016; Betti *et al*., 2016).

The CBC has been subject to continued massive selection due to changing atmospheric CO_2_ and O_2_ levels, and probably also in response to changing temperature and water and nutrient supply (Sage, 2017; Raven *et al*., 2017). This included large changes in the structure and characteristics of Rubisco, with plant Rubiscos having a higher selectivity for CO_2_ over O_2_, but a lower *k*_*cat*_ than those of ancestral algae (Jordan & Ogren, 1981; Badger *et al*., 1998; Savir *et al*., 2010; Shih *et al*., 2016; Sharwood *et al*., 2016; Erb & Zarzycki, 2018; Iniguez *et al*., 2020). Higher selectivity for CO_2_ may be mechanistically linked to decreased *k*_*cat*_ (Tcherkez *et al*., 2006; Young *et al*., 2016; Flamholz *et al*., 2019). As Rubisco is the most abundant protein in plant leaves (Ellis, 1979) and on the globe (Bar-On & Milo, 2019) this trade-off impacts photosynthetic nitrogen use efficiency at the plant and ecosystem level. Little is known about whether evolutionary pressures drove other changes in the CBC (Stitt *et al*., 2021).

Several globally-important groups of photosynthetic organisms have evolved inorganic carbon-concentrating mechanisms (CCMs) that concentrate CO_2_ and suppress the wasteful side reaction of Rubisco with O_2_. The earliest appearance of cyanobacterial CCM is controversial (400 – 2000 million years ago; Badger & Price, 2003; Kupriyanova *et al*., 2013). The well conserved beta-carboxysomes likely evolved quite early before the large cyanobacterial radiation occurred (Melnicki *et al*., 2021). The origin of algal CCMs has been related to the appearance of equimolar CO_2_:O_2_ concentrations in surface waters around 500 million years ago (Griffiths *et al*., 2017). A CCM termed C_4_ photosynthesis evolved 25-30 million years ago in some terrestrial angiosperms, coinciding with a transition in the Earth’s climate from hot and wet conditions with atmospheric CO_2_ concentrations >1000 ppm to cooler and drier conditions and CO_2_ concentrations <300 ppm (Zachos *et al*., 2008).

Cyanobacteria evolved a so-called biophysical CCM, which aims to accumulate high amounts bicarbonate in the cell. This ion can be actively transported and cannot easily escape through membranes from the cell. They use three bicarbonate transporters and two CO_2_ hydrating/uptake systems to actively concentrate bicarbonate within the cells. Bicarbonate then diffuses into the bacterial microcompartment carboxysome, where it is converted to CO_2_ by carbonic anhydrase (CA) to generate a high concentration of CO_2_ around Rubisco (Kaplan & Reinhold, 1999; Rae *et al*., 2013; Hagemann *et al*., 2021). Carboxysomes are composed of about ten proteins including a self-assembling proteinaceous sheath that encapsulates Rubisco and CA as well as proteins that are required for assembly and organisation of Rubisco (Price *et al*., 2013; Kerfeld & Melnicki, 2016; Kerfeld *et al*., 2018; Lechno-Yossef *et al*., 2020). Mutations that abolish bicarbonate accumulation or carboxysome function lead to loss of the ability to grow at ambient CO_2_ concentrations, but can be rescued at enhanced CO_2_ levels (Badger *et al*., 1991; Marcus *et al*., 1992; So *et al*., 2002; Xu *et al*., 2008).

Many eukaryotic algae evolved an alternative biophysical CCM that accumulates CO_2_ around Rubisco in the pyrenoid (Badger *et al*., 1998; Borkhsenious *et al*., 1998; Giordano *et al*., 2005; Wang *et al*., 2015; Meyer *et al*., 2017; Griffiths *et al*., 2017). The pyrenoid is a dynamic liquid-like microstructure (Freeman Rosenzweig *et al*., 2017; Atkinson *et al*., 2020; Barret *et al*., 2021) containing over 100 proteins (MacKinder *et al*., 2017; Zhan *et al*., 2018; Küken *et al*., 2018) including Rubisco and Rubisco activase. Rubisco is recruited to the pyrenoid matrix by the protein EPYC1 (MacKinder *et al*., 2016; Atkinson *et al*., 2019; Meyer *et al*., 2020). This process depends on Rubisco small subunit (Meyer *et al*., 2012) and the SAGA1 protein (Itakura *et al*., 2019), which also plays a role in assembling the surrounding starch sheath. The pyrenoid is penetrated by a dense network of tubules deriving from the thylakoid (Engel *et al*., 2015). Bicarbonate transporters located on the plasma membrane and plastid envelope membrane (Karlsson *et al*., 1998; Wang *et al*., 2014; Yamano *et al*., 2015) and bestrophin-like proteins in the thylakoid (Mukherjee *et al*., 2019) concentrate bicarbonate into the thylakoid lumen where it is converted to CO_2_ by the carbonic anhydrase CAH3 (Karlsson *et al*., 1998; Mitra *et al*., 2005; Sinetova *et al*., 2012; Blanco-Rivero *et al*., 2012), with the acidic lumen pH driving the reaction towards CO_2_ formation (Moroney & Ynalvez, 2007; Spalding, 2008; Burlacot *et al*., 2022). Loss of bicarbonate or CO_2_ transporters, misexpression of carbonic anhydrase or disturbed pyrenoid structure leads to attenuation or loss of the ability to grow at ambient CO_2_ (Griffiths *et al*., 2017; Caspari *et al*., 2017; Toyokawa *et al*., 2020).

In C_4_ plants, Rubisco and the remainder of the CBC are restricted to bundle sheath cells located around the vasculature in the centre of the leaf. This biochemical CCM initially fixes inorganic carbon as bicarbonate by phospho*enol*pyruvate (PEP) carboxylase in the mesophyll to generate 4-C metabolites that move to the bundle sheath where they are decarboxylated to release CO_2_ and 3-C metabolites, which move back to the mesophyll (Hatch, 2002; von Caemmerer & Furbank, 2003; Sage *et al*., 2012; Sage, 2017; Arrivault *et al*., 2019).

CCMs impact on operation of the CBC. One well characterized impact is that high CO_2_ concentrations allow Rubisco to keep (cyanobacteria and algae) or to revert (C_4_ plants) to a lower-selectivity and higher *k*_*cat*_ form (Pearce, 2006; Tcherkez *et al*., 2006) with an associated decrease in the amount of nitrogen that must be invested in Rubisco. This has been well-documented in cyanobacteria (Andrews & Abel, 1981; Badger & Price, 1992; Read, 1994; Kerfeld & Melnicki, 2016), pyrenoidal green algae (Badger *et al*., 1998; Kaplan & Reinhold, 1999; Meyer & Griffiths, 2013; Heureux *et al*., 2017; Goudet *et al*., 2020) and terrestrial C_4_ plants (Brown, 1978; Yeoh *et al*., 1980; Sharwood *et al*., 2016; Sage, 2017). Little is known about whether CCMs require further modifications of the CBC. However, this is plausible. For example, concentration of CO_2_ is associated with increased energy demand in cyanobacteria (Kaplan *et al*., 1987; Marcus *et al*., 1992; Badger *et al*., 2006; Price *et al*., 2013), eukaryotic algae (Badger *et al*., 1998; Meyer *et al*., 2017; Griffiths *et al*., 2017; Burlacot *et al*., 2022) and C_4_ photosynthesis (von Caemmerer & Furbank, 2003; Zhu *et al*., 2008; Sage, 2017; Bräutigam *et al*., 2018). This additional energy demand may affect poising between photosynthetic electron transport and C assimilation by the CBC (Burlacot *et al*., 2022). Furthermore, cyanobacterial and algal CCMs require spatial separation of Rubisco from the remainder of the CBC. Rubisco is located inside and all other CBC enzymes are located outside the carboxysome (Price *et al*., 2013; Kerfeld & Melnicki, 2016; Kerfeld *et al*., 2018; Sun *et al*., 2019). A similar picture holds for the pyrenoid in *C. reinhardtii* (MacKinder *et al*., 2016; Küken *et al*., 2018; Zhan *et al*., 2018). This raises questions on how RuBP and 3PGA move between spatially-separated enzymes.

Metabolite profiling provides an unbiased strategy to search for inter-species variance in pathway operation (Stitt *et al*., 2021); changes in the balance between different enzymatic steps will lead to changes in the relative levels of pathway intermediates, irrespective of whether the underlying cause involves changes in protein abundance, enzyme kinetics, enzyme regulation or changes in the high-level structure of regulatory networks. Information about the levels of CBC metabolites is rather sparse. Whilst there were studies of CBC metabolite levels in various plant species in the 1980s (summarised in Stitt *et al*., 2021) for technical reasons they focused a handful of metabolites (RuBP, 3PGA, triose-P, fructose 1,6-bisphosphate (FBP), fructose-6-phosphate (F6P)). Similar restricted analyses have been performed for CBC intermediates in cyanobacteria (Marcus *et al*., 2011). Extensive profiling of CBC metabolites was performed by Bassham and colleagues in *Chlorella pyrenoidosa* (Bassham & Kirk, 1960; Pedersen *et al*., 1966; Bassham & Krause, 1969) using steady-state ^14^CO_2_ labelling followed by 2-dimensional paper chromatography and autoradiography. However, this approach is not applicable to higher plants (Stitt *et al*., 2021).

Analytic platforms that combine chromatographic separation with tandem mass spectrometry (LC-MS/MS) allow systematic profiling of CBC metabolites (Cruz *et al*., 2007; Arrivault *et al*., 2009; Hasunuma *et al*. 2010; Ma *et al*., 2014). They have recently been used to compare CBC metabolite profiles in several C_3_ and C_4_ species, and in species that represent steps in the evolution of C_4_ photosynthesis (Arrivault *et al*., 2019; Borghi *et al*., 2019; Borghi *et al*., 2021; Stitt *et al*., 2021). This comparative approach revealed variation in CBC metabolite profiles between different C_4_ species and between different C_3_ species. Despite this variance within each photosynthesis-type, the differences in CBC metabolite profile were large enough to clearly separate C_4_ species from C_3_ species (Arrivault *et al*., 2019; Borghi *et al*., 2021). Separation was partly driven by RuBP being lower in C_4_ species than in C_3_ species. Much of the RuBP is bound in the active site of Rubisco (von Caemmerer & Farquhar, 1985; Woodrow & Berry, 1988; Sage *et al*., 1988; Salvucci, 1989; Parry *et al*., 2008; Sharkey, 2022). A plausible explanation for the lower RuBP in C_4_ species is that they have a lower abundance of Rubisco protein than C_3_ species (see above).

In view of their large contribution to global C exchange (see above) and their importance in biotechnology (Doron *et al*., 2017), it is important to learn more about how the CBC operates in cyanobacteria and algae. To our knowledge, CBC metabolite levels in cyanobacteria and eukaryotic algae have not been systematically profiled, except for studies of *Chlamydomonas reinhardtii* under high CO_2_ (Mettler *et al*., 2014; Küken *et al*., 2018) and Rubisco-deficient *C. reinhardtii* mutants in mixotrophic conditions (Saint-Sorny *et al*., 2022).

Here, we profile CBC metabolites under ambient CO_2_ concentration in one cyanobacterial species (*Synechocystis* sp. PCC 6803, hereafter *Synechocystis*) and three eukaryotic algae: *C. reinhardtii, Chlorella sorokiniana and Chlorella ohadii. Synechocystis* has long served as a model for analysis of photosynthesis in cyanobacteria (Ikeuchi & Tabata, 2001). The three eukaryotic algae derive from two different phylogenetic branches of the chlorophyte algae. *C. reinhardtii* has long been a model for photosynthesis research (Rochaix, 2002; Merchant *et al*., 2007; Salome & Merchant, 2017), whilst the recently-discovered *C. ohadii* (Treves *et al*., 2013) is attracting attention because it is the fastest growing algae characterised to date and possesses remarkable ability to withstand stress (Treves *et al*., 2017; 2020; 2022). We then combine these cyanobacterial and algal metabolite profiles with recently published profiles from terrestrial plant species to ask i) if CBC metabolite profiles differ enough to distinguish cyanobacteria and algae from C_3_ and C_4_ terrestrial plants and ii) if their CCMs impact on CBC operation in a similar or different way to the impact of the CCM in C_4_ photosynthesis.

## Materials and Methods

*Synechocystis* was cultured in glass tubes (Rippka *et al*., 1979) and algae in flat glass bioreactors (Treves *et al*., 2022). Cultures were bubbled with air, illuminated at 100 µmol photons m^-2^ s^-1^ and harvested under illumination in cold (−70°C) 70% methanol. This is essential due to the very rapid turnover of CBC metabolites 0.1-1 sec, (Stitt *et al*., 1980; Arrivault *et al*., 2009). Metabolites were extracted following Mettler *et al*. (2014), spiked with stable-isotope-labelled standards (Arrivault *et al*., 2015) and quantified by using reverse phase (Arrivault *et al*., 2009) or (3PGA) anion exchange LC-MS/MS (Lunn *et al*., 2006). 3PGA was also quantified enzymatically (Merlo *et al*., 1993). For details see Method S1.

## Results

### CBC metabolite profiles

The cyanobacterium *Synechocystis* and the three green algae *C. reinhardtii, C. sorokiniana* and *C. ohadii* were cultured in CCM-inducing conditions (ambient CO_2_) at low irradiance (100 µmol photons m^-2^ s^-1^) and harvested by rapid injection under continued illumination into very cold 70% methanol to profile CBC metabolite levels. Quantification was supported by spiking with stable-isotope labelled standards. Metabolite levels were initially expressed on a dry weight (DW) basis (Dataset S1).

In all cases, the profile was dominated by high RuBP and 3PGA, whereas the levels of all other metabolites were relatively low (Fig. 1). Within the RuBP regeneration sector, sedoheptulose 1,7-bisphosphate (SBP) and sedoheptulose-7-phosphate (S7P) were much higher than the other metabolites in *Synechocystis* and somewhat higher in *C. reinhardtii*, whereas they were similar to or lower than FBP and F6P in *C. sorokiniana* and *C. ohadii*. The various pentose phosphates (ribose-5-phosphate (R5P) and ribulose-5-phosphate + xylulose-5-phosphate (Ru5P+Xu5P)) were low in all four species. Quantification of 3PGA was complicated by ion suppression when analysed by reverse phase LC-MS/MS. The samples were therefore also analysed by anion exchange LC-MS/MS and by enzymatic assay. The latter approach gave lower results, except for *C. reinhardtii* (see Table S1). Based on these results and the fact that quantification by enzymatic assay was not trivial, 3PGA was quantified by anion exchange LC-MS/MS. Glyceraldehyde-3-phosphate (GAP) was below the level of detection of our LC-MS/MS platforms. A large excess of dihydroxyacetone phosphate (DHAP) over GAP is expected, based on the equilibrium constant of the triose phosphate isomerase reaction. Much higher levels of GAP than DHAP were previously reported after enzymatic assay in *Synechocystis* (Marcus *et al*., 2011). We therefore reanalysed our *Synechocystis* samples with and without spiked authentic GAP standard. The signal for GAP in unspiked samples was at or below the level of detection (Fig. S1).

**Fig. 1.**
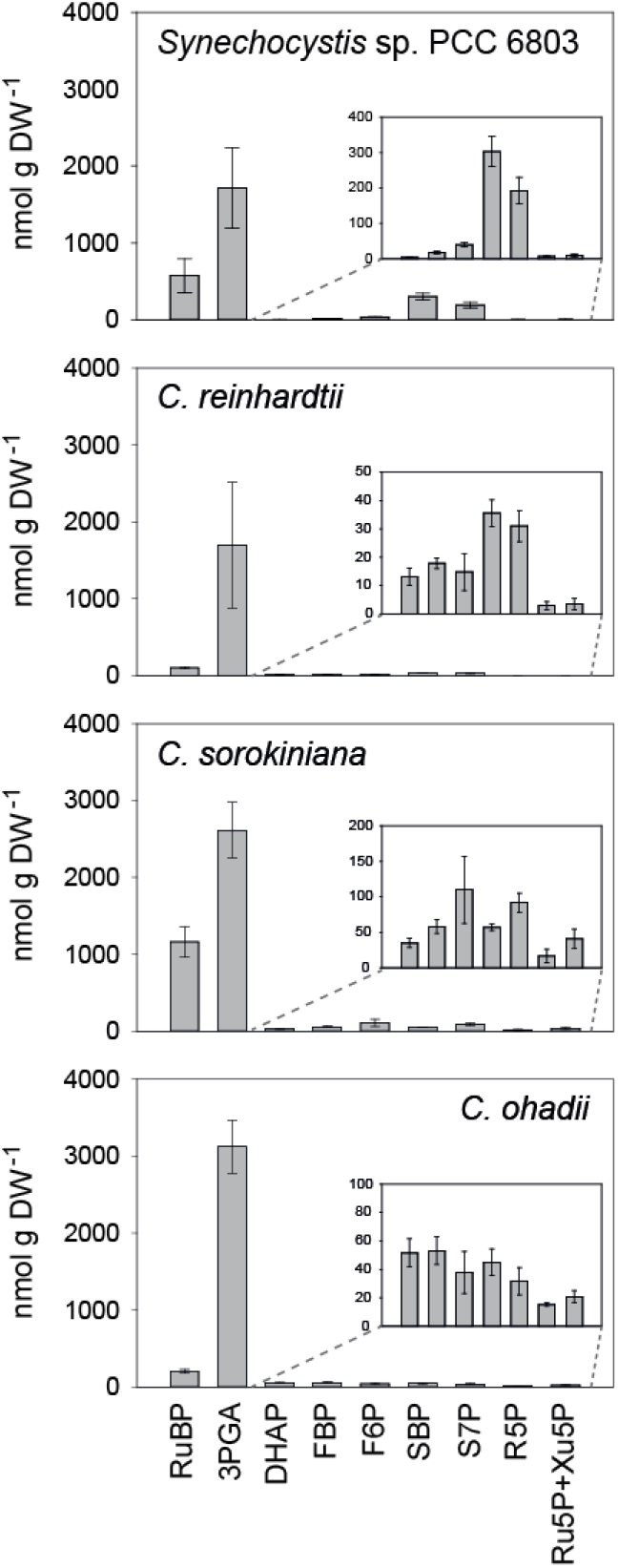
CBC metabolite profiles in cyanobacteria and algae. *Synechocystis* sp. PCC 6803, *Chlamydomonas reinhardtii, Chlorella sorokiniana and Chlorella ohadii* were incubated at 100 µmol quanta m^-2^ s^-1^ and rapidly quenched into cold methanol. The inserts show, with an expanded y-axis, the levels of DHAP and metabolites in the RuBP regeneration sector of the CBC. The results are shown as mean (nmol g FW^-1^) ± SD (*n*=3 to 4). Data are presented in Dataset S1.

We compared the cyanobacterial and algal CBC metabolite profiles with published data for a large set of terrestrial plant species including seven C_3_ species (*Arabidopsis thaliana, Manihot esculenta, Nicotiana tabacum, Triticum aestivum, Oryza sativa, Flaveria robusta, Flaveria cronquistii*), two *Flaveria* species that represent intermediate steps in the evolution of C_4_ photosynthesis (*Flaveria ramosissima, Flaveria anomala*), three C_4_-like species (*Flaveria palmieri, Flaveria vaginata, Flaveria brownii*) and four C_4_ species (*Flaveria bidentis, Flaveria trinervia, Zea mays, Setaria viridis)* (Arrivault *et al*, 2019; Borghi *et al*., 2021; summarised in Fig. S2, data presented in Dataset S1). In all cases, leaf material was harvested in growth conditions under moderately limiting irradiance. For terrestrial plant species, 3PGA was analysed enzymatically and metabolite levels expressed on a fresh weight (FW) basis. Visual inspection revealed that the profiles in cyanobacteria and algae were very distinctive from those in terrestrial plants. In particular, terrestrial plant species did not contain such a marked excess of both RuBP and 3PGA compared to other CBC metabolites (Fig. S2). The excess of SBP and S7P over other metabolites in the RuBP regeneration sector of the cycle in *Synechocystis* was also not seen in any terrestrial species. The relative levels of CBC metabolites in *Z. mays, A. thaliana* and *O. sativa* leaves do not change greatly over a wide range of limiting irradiance (Arrivault *et al*., 2019; Borghi *et al*., 2019). Moreover, similar trends were also observed in *C. ohadii* cells grown and harvested at ambient CO_2_ under 3000 µmol photons m^-2^ s^-1^ (Treves *et al*., 2022). It is therefore unlikely that the very high RuBP and 3PGA in cyanobacteria and algae can be explained by differences in irradiance.

### Principal components analysis of CBC intermediate profiles separates cyanobacteria and green algae from plant species

Principal components (PC) analysis was performed to provide a systematic comparison of CBC metabolite profiles in cyanobacteria, green algae and terrestrial plant species. The cellular composition of these plant species is very different in terms of protein and chlorophyll content (Arrivault *et al*., 2019). Furthermore, metabolite amounts were not expressed on the same basis for the species studied (on DW for cyanobacteria and algae, on FW for terrestrial plant species). To avoid bias due to differences in cellular composition between species, we normalized metabolite levels by expressing the C in each metabolite as a fraction of the total C in all CBC metabolites in that species (for calculation, see Dataset S1, Method S1). The normalized data set provides information about the balance between different reactions in the CBC, as captured by the distribution of C between metabolites in the pathway (see Arrivault *et al*., 2019; Stitt *et al*., 2021).

PC1 (29%), PC2 (25%) and PC3 (17%) captured over 70% of total variance, with further PCs making only a small contribution (see Table S2). Cyanobacteria and algae were clearly separated from terrestrial plant species in the PC1 versus PC2 (Fig. 2A) and the PC1 versus PC3 (Fig. 2B) plot. *Synechocystis* was more closely related to *C. sorokiniana* than to *C. reinhardtii* and *C. ohadii*. Among the plant species, C_4_ species were separated from C_3_ species, with *Flaveria* C_3_-C_4_ intermediate species and C_4_-like species lying between *Flaveria* C_3_ species and *Flaveria* C_4_ species, as previously seen when PC analysis was performed without cyanobacteria and algae (Borghi *et al*., 2021; Supplemental Fig. S3). The cyanobacteria and algae separated orthogonally to the C_4_ species, even though all these organisms operate a CCM (however, biophysical versus biochemical, respectively). Inspection of loadings revealed that the separation of cyanobacteria and algae from plant species was driven by 3PGA, RuBP and (for *Synechocystis*) high SBP in the PC1 versus PC2 analysis, and by high 3PGA and RuBP in the PC1 versus PC3 analysis.

**Fig. 2.**
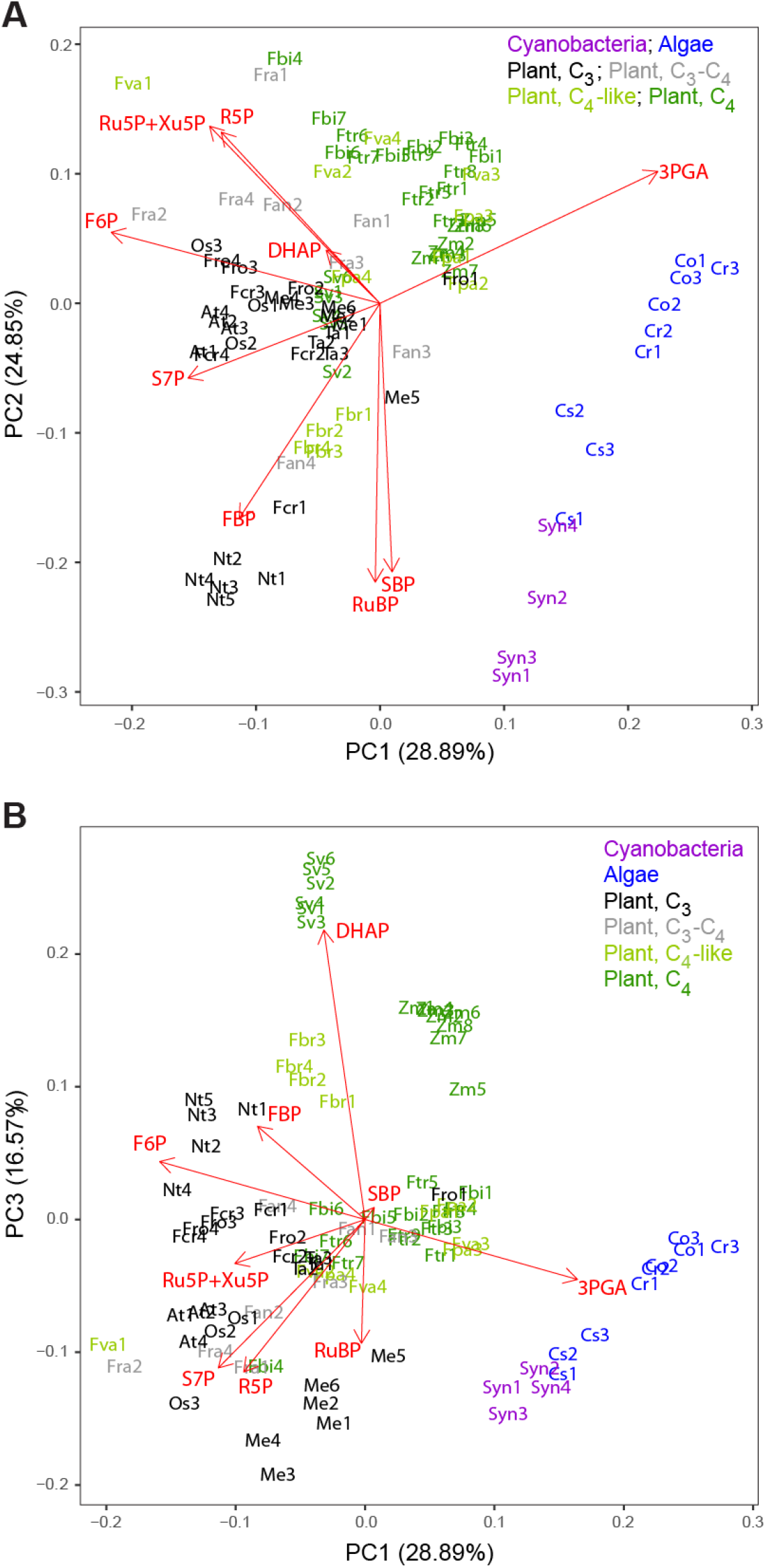
Principal components analysis of CBC metabolite profiles in cyanobacteria, algae and terrestrial plant species. The PC analysis was performed after normalising the amount of C in a given metabolite on the total amount of C in all CBC metabolites in that species. The contribution of PC1 to PC10 to total variance is shown in Table S2. **(A)** PC1 versus PC2, **(B)** PC1 versus PC3. Cyanobacteria, algae, C_3_, C_4_, C_3_-C_4_ intermediate and C_4_-like terrestrial plants are denoted by colour (purple, blue, black, grey, pale green and green, respectively; see insert legend). Abbreviations: Species abbreviations, full name and photosynthesis mode are, alphabetically: At, *Arabidopsis thaliana* (C_3_); Cr, *Chlamydomonas reinhardtii* (pyrenoidal alga); Fan, *Flaveria anomala* (C_3_-C_4_); Fbi, *Flaveria bidentis* (C_4_); Fbr, *Flaveria brownii* (C_4_-like); Fcr, *Flaveria cronquistii* (C_3_); Fpa, *Flaveria palmeri* (C_4_-like); Fra, *Flaveria ramosissima* (C_3_-C_4_); Fro, *Flaveria robusta* (C_3_); Ftr, *Flaveria trinervia* (C_4_); Fva, *Flaveria vaginata* (C_4_-like); Me, *Manihot esculenta* (C_3_); Nt, *Nicotiana tabacum* (C_3_); Co, *Chlorella ohadii* (pyrenoidal alga); Os, *Oryza sativa* (C_3_); Cs, *Chlorella sorokiniana* (pyrenoidal alga); Sv, *Setaria viridis* (C_4_); Syn, *Synechocystis* sp. PCC 6803 (cyanobacteria with carboxysome); Ta, *Triticum aestivum* (C_3_); Zm, *Zea mays* (C_4_). Normalized data are presented in Dataset S1, including terrestrial plant data taken from Arrivault *et al*. (2019) and Borghi *et al*. (2021).

### Cross-species comparison of individual metabolite levels

To provide more information on inter-species differences in CBC metabolites, we compared their individual levels between cyanobacteria, algae and the various terrestrial plant species (Fig. 3). Metabolite levels were again expressed as % of total C in all CBC metabolites.

**Fig. 3.**
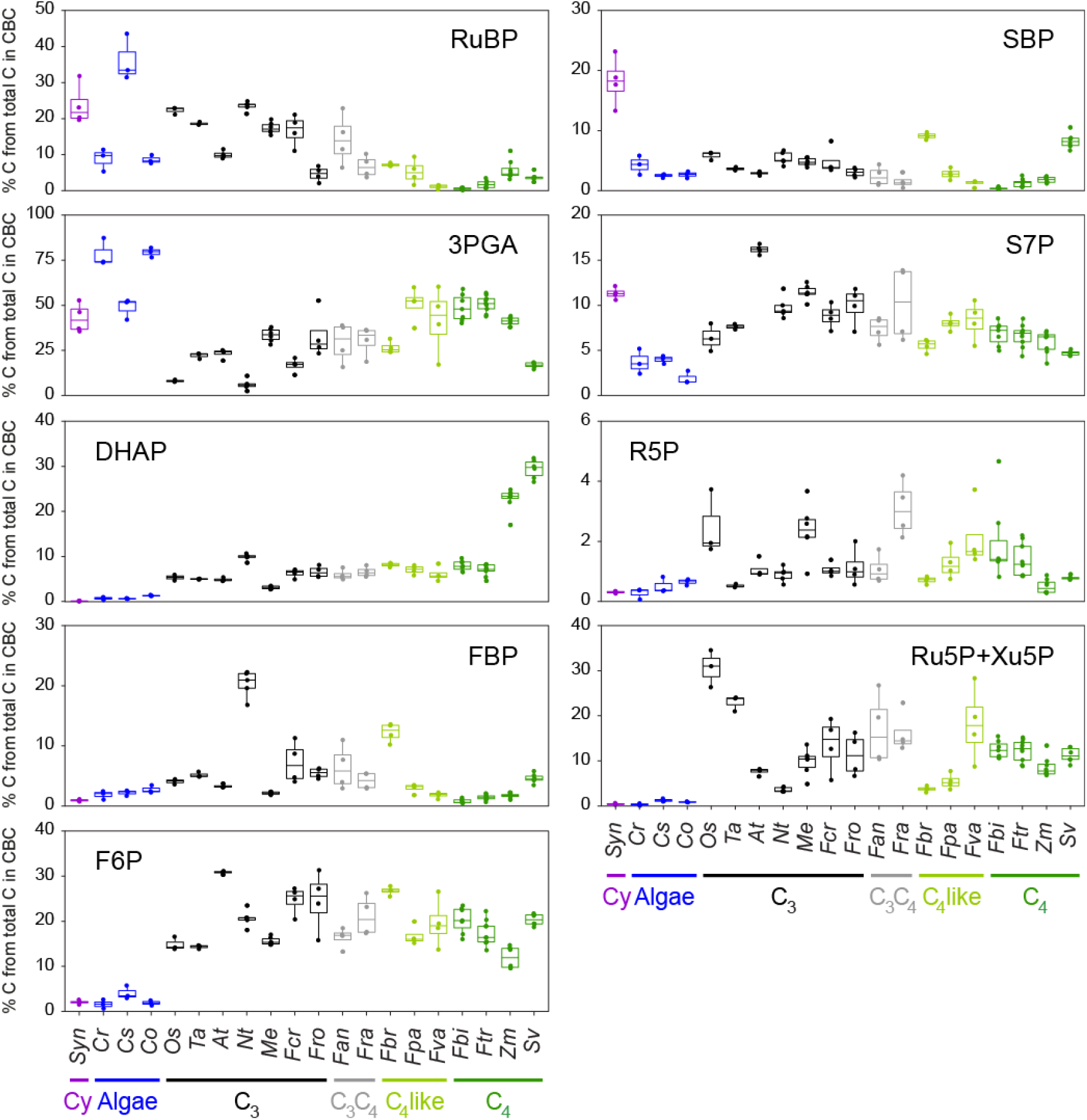
Cross-species comparisons of metabolite levels. The amount of C in a given metabolite was normalised on the total amount of C in all CBC metabolites in that species. The data are displayed as box plots (*n*=3 to 9). Species are ordered from left to right, cyanobacteria (Cy, purple), algae (blue), C_3_ (black), C_3_-C_4_ intermediate (grey), C_4_-like (light green) and C_4_ terrestrial plant species (green). For species abbreviations see legend of Fig. 2. Normalized data are presented in Dataset S1, including terrestrial plant data taken from Arrivault *et al*. (2019) and Borghi *et al*. (2021).

Comparing first across the reference set of terrestrial plants, there were no consistent differences in CBC metabolite allocation based on photosynthesis mode except for RuBP, 3PGA and, to a lesser extent, DHAP. Allocation to RuBP decreased and allocation to 3PGA increased along the axis from C_3_ to C_3_-C_4_ intermediate, C_4_-like and terrestrial plant C_4_ species. A slight increase was observed for DHAP along this axis, with especially high values in the C_4_ species *Z. mays* and *S. viridis*. The downwards trend in RuBP reflects the increasing *k*_*cat*_ of Rubisco and the resulting decrease in the abundance of Rubisco and hence RuBP-binding sites (see Introduction). The high 3PGA and DHAP in C_4_ plants reflects the high levels of these metabolites that are required to support the concentration gradients which drive their movement in an intercellular shuttle that transfers NADPH and ATP from the mesophyll to bundle sheath cells in NADP-ME type C_4_ species (Leegood, 1985; Stitt & Heldt, 1985; Arrivault *et al*., 2017). However, values in C_4_ Flaveria species were similar to C_3_-C_4_ intermediate and C_4_-like species.

Cyanobacteria and algae were characterized by high allocation to both RuBP (similar or higher than C_3_ species and much higher than in C_4_ species) and 3PGA (much higher than in C_3_ species and similar or higher than C_4_ species). They exhibited lower levels of DHAP (especially in S*ynechocystis*), F6P, S7P (except *Synechocystis*), R5P and Ru5P+Xu5P than terrestrial plant species, irrespective of their photosynthesis modes. Allocation to FBP and SBP in cyanobacteria and algae fell within the range observed in terrestrial plants, with the exception that SBP was higher in *Synechocystis*.

### Cross-species comparison of metabolite ratios

Metabolite ratios provide an indicator for the balance between different reactions in a pathway. The RuBP/3PGA provides an indicator for restriction at the carboxylation reaction of Rubisco, the 3PGA/DHAP ratio for the supply of NADPH and ATP (viz. the balance between energy production in the light reactions and energy consumption in the CBC, Dietz & Heber, 1984), the FBP/F6P and SBP/S7P ratios are indicators for resistance to flux at fructose-1,6-bisphosphatase (FBPase) and sedoheptulose-1,7-bisphosphatase (SBPase), respectively, and the pentose-P/RuBP ratio for resistance to flux at phosphoribulokinase (PRK). In the latter case, R5P and Ru5P+Xu5P were summed as pentose-P. Ratios are shown for cyanobacterial, algal and terrestrial plant species in Fig. 4.

**Fig. 4.**
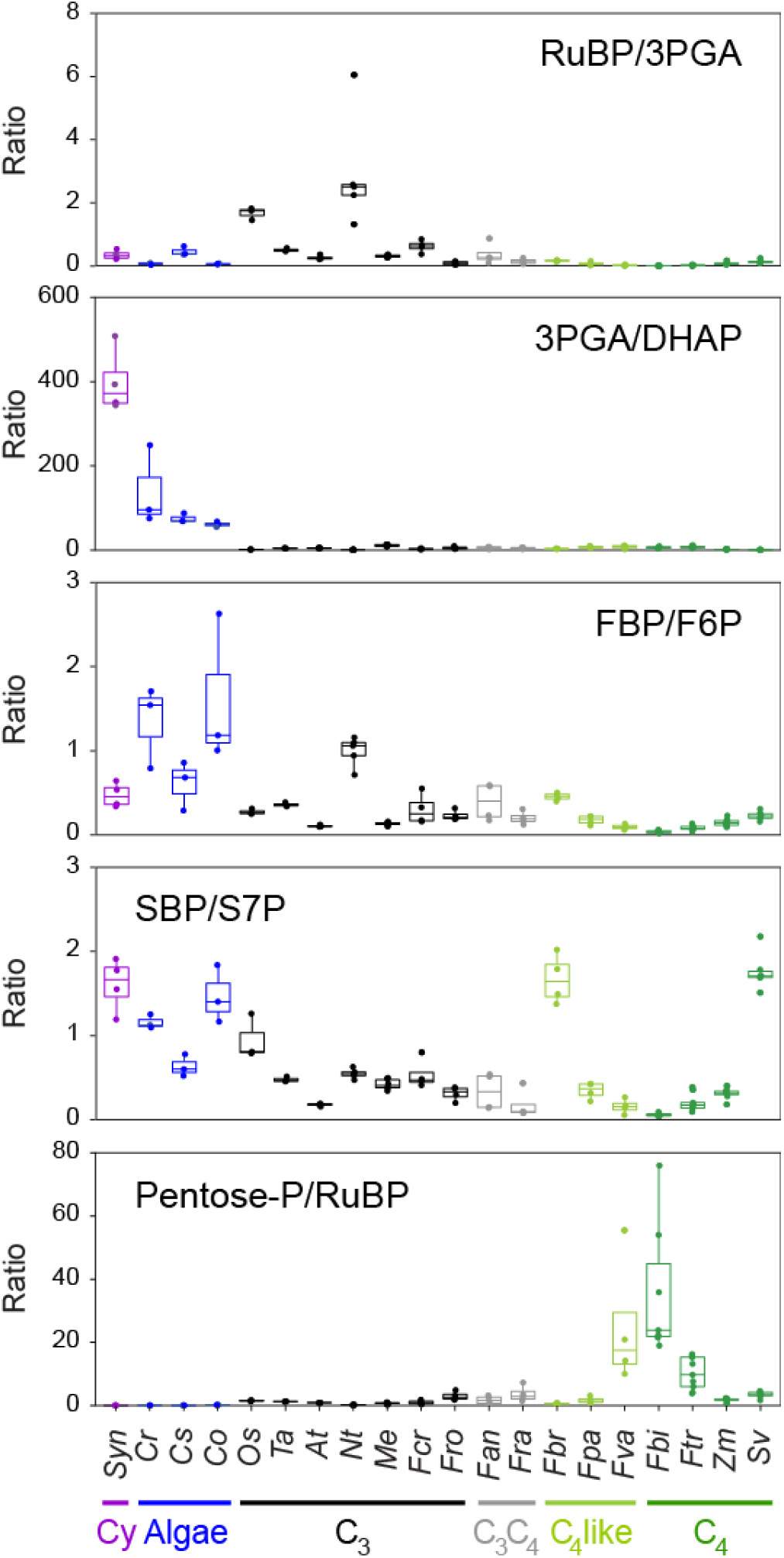
Cross-species comparison of metabolite ratios. Ratios were determined using metabolite amounts. The data are displayed as box plots (*n*=3 to 9). Species are ordered from left to right, cyanobacteria (Cy, purple), algae (blue), C_3_ (black), C_3_-C_4_ intermediate (grey), C_4_-like (light green) and C_4_ terrestrial plant species (green). For species abbreviations see legend of Fig. 2. For separated plots of metabolite ratios in cyanobacteria, algae and in terrestrial plant species see Fig. S4. Ratios are presented in Dataset S1, including terrestrial plant data taken from Arrivault *et al*. (2019) and Borghi *et al*. (2021).

We observed a clear trend to lower RuBP/3PGA ratios in cyanobacteria and algae than in plant C_3_ species, but no consistent difference between them and plant species that operate a CCM, i.e., C_3_-C_4_ intermediate, C_4_-like and C_4_ species. The 3PGA/DHAP ratio was far higher in cyanobacteria and algae than in any of the terrestrial plant species, reflecting the very high 3PGA and relatively low DHAP in cyanobacteria and algae (see Discussion). Their FBP/F6P and SBP/S7P ratios tended to be higher than in plant species, with the exception of the FBP/F6P ratio for *N. tabacum*, and the SBP/S7P for *O. sativa, F. brownie and S. viridis* which were similar or higher than for cyanobacteria and algae. This points to a larger restriction of RuBP regeneration by FBPase and SBPase in cyanobacteria and algae than in many plant species. The pentose-P/RuBP ratio was much lower in cyanobacteria and algae than in the plant species. This reflects the low pentose-P and high RuBP levels in cyanobacteria and algae, and indicates upregulation of PRK.

In many cases, metabolite ratios were so different between cyanobacteria, algae and plants that displaying them on a single scale masked differences between cyanobacteria and algae or between plant species. In Fig. S4, metabolite ratios are plotted using different scales for cyanobacteria and algae and for terrestrial plant species. Among cyanobacteria and algae, the RuBP/3PGA ratio was higher in S*ynechocystis* and *C. sorokiniana* than in *C. reinhardtii* and *C. ohadii*, and the 3PGA/DHAP ratio was higher in *Synechocystis* than in the three eukaryotic algae. The plant species showed a trend to a decreased RuBP/3PGA and FBP/F6P ratio and increased pentose-P/RuBP ratio along the axis from C_3_ to C_3_-C_4_ intermediate, C_4_-like and C_4_ species.

We also compared the ratios of DHAP/FBP, DHAP/F6P, DHAP/SBP, DHAP/S7P and DHAP/pentose-P across all species (Fig. S5). DHAP is the starting point for regeneration of RuBP. A higher ratio of DHAP to these sugar-phosphates would indicate increased resistance to flux at various sites in the regenerative phase of the CBC. In general, the ratios in cyanobacteria and algae were very low due to the small DHAP pool in these species. Hence, their DHAP/FBP and DHAP/SBP ratios were lower than in any plant species, irrespective of photosynthesis mode. These very low ratios may point to decreased resistance to flux at aldolase. It is also possible that they reflect a lower contribution of a cytosolic DHAP pool in green algae, compared to plants where over half of the DHAP is in the cytosol (Stitt *et al*., 1980; Gerhardt *et al*., 1987). The extremely low DHAP/FBP and DHAP/SBP ratios in S*ynechocystis* reflect the very low DHAP in this species (see Fig. 3), which may be partly due to the absence of subcellular compartmentation in cyanobacteria.

In terrestrial plant species, there was a trend to an increase of all ratios along the axis from C_3_ to C_3_-C_4_ intermediate, C_4_-like and C_4_ species (with the exception of the DHAP/SBP ratio for *S. viridis*). As already mentioned, these high ratios reflect the presence of a large DHAP pool in the mesophyll cells to drive diffusion back to the bundle sheath cells in these NADP-ME type C_4_ species rather than a change in poising within the CBC itself. Incidentally, these high values were absent for DHAP/F6P, DHAP/S7P and DHAP/pentose-P ratios in C_4_ *Flaveria* species and for all ratios in the C_4_-like *Flaveria* species, indicating that optimisation of the energy shuttle was one of the events that occurred late during the evolution of C_4_ photosynthesis.

## Discussion

Cyanobacteria and eukaryotic algae play a major role in the global C cycle (Mann, 1996; Field *et al*., 1998; Behrenfeld *et al*., 2001; Thierstein & Young, 2004; Meyer & Griffith, 2013; Rousseau & Gregg, 2013). Due to the low and often fluctuating concentrations of CO_2_ in water at the slightly alkaline pH that reigns in most marine and freshwater environments, their photosynthesis depends on the operation of a CCM that allows intracellular accumulation of bicarbonate and its use to generate high CO_2_ concentrations in a microcompartment that contains Rubisco (see Introduction). Cyanobacteria and algae contain fundamentally different microcompartments, termed carboxysomes and pyrenoids, respectively, but they share in common that Rubisco is physically separated from the remainder of the CBC enzymes, which are located in the free cytoplasm of cyanobacteria (Price *et al*., 2013; Kerfeld & Melnicki, 2016; Kerfeld *et al*., 2018) and in the chloroplast stroma of eukaryotic algae (MacKinder *et al*., 2016; Zhan *et al*., 2018; Küken *et al*., 2018). For each molecule of CO_2_ that is assimilated, one molecule of RuBP will need to move into and two molecules of 3PGA will need to move out of the carboxysome or pyrenoid. It can be assumed that the conductivity of the surface of these microcompartments for small molecules is kept relatively low, because otherwise there would be rapid back leakage of bicarbonate and CO_2_, resulting in a futile cycle that would waste considerable amounts of energy. This raises questions about how RuBP and 3PGA move into and out of these microcompartments.

The cyanobacterial beta-type carboxysome found in *Synechocystis* is surrounded by a proteinaceous shell that is mainly composed by three shell protein types. Most abundant are hexameric CcmKs forming a small central pore that is rich in positively charged amino acid residues, which is believed to be the main gate for bicarbonate entry by diffusion but probably not permitting CO_2_ and O_2_ diffusion (Liu, 2022). The exchange of the CBC intermediates 3PGA and RuBP is likely achieved by CcmO and CcmP, two less abundant shell proteins that form larger central pores. Especially CcmP is discussed as a gate for 3PGA and RuBP diffusion, since crystallographic studies showed signs for bound 3PGA (Cai *et al*., 2013) or possibly RuBP (Larsson *et al*., 2017) in its central pore, which is also able to form open or closed conformations permitting regulation of transport (reviewed in Liu, 2022). Less is known about the molecular details of the pyrenoid surface, except that a starch sheath probably reduces back-leakage of CO_2_ (Toyokawa *et al*., 2020). This sheath will presumably also hinder movement of RuBP and 3PGA.

We have shown that the CBC metabolite profiles in a cyanobacteria and three eukaryotic algae differ profoundly from those in a large set of terrestrial plant species, including C_4_ species (Fig. 2, compare also Fig.1 and Fig. S2). In particular, cyanobacteria and algae contained high levels of RuBP and particularly 3PGA compared to the levels found in plants (Fig. 3). The high abundance of RuBP was less marked compared to C_3_ species but was striking compared to C_4_ species, which also operate a CCM. The relatively high RuBP in C_3_ plants may reflect the high abundance of Rubisco, which will result in a large amount of RuBP being at Rubisco active site (see Introduction and below). The high abundance of 3PGA was marked compared to C_3_ species, and less so compared to some C_4_ species where 3PGA has an additional ancillary role in a reductant/energy shuttle that requires a large 3PGA pool to drive rapid diffusion from bundle sheath to mesophyll cells (Leegood, 1985; Stitt & Heldt, 1985).

The most obvious explanation for the large pools of RuBP and 3PGA in cyanobacteria and algae is that they are required to generate concentration gradients that drive movement of RuBP into the carboxysome or pyrenoid, and movement of 3PGA out of the carboxysome or pyrenoid. Several other features of the CBC metabolite profiles in S*ynechocystis* and these algae species are consistent with this proposal. One is that there was always a clear excess of 3PGA over RuBP, ranging from a three-fold excess in S*ynechocystis* and *C. sorokiniana* to a ten-fold excess in *C. reinhardtii* and *C. ohadii*. This matches the expectation from reaction stoichiometry (see above) that 3PGA must move twice as quickly as RuBP. Another feature was the remarkably high 3PGA/DHAP ratios in algae, which lay in the range of 80-400, compared to 1-10 in plants (Fig. 4, Fig. S4). Whilst it is in principle possible that this reflects a massive imbalance between the production of ATP and NADPH in the light reactions and consumption of NADPH and ATP in the CBC and the CCM, the most likely explanation for the high 3PGA/DHAP ratio in cyanobacteria and algae is that much of the 3PGA is located inside the carboxysome or pyrenoid, and that the amount of 3PGA that is available for reduction by phosphoglycerate kinase and NADP-glyceraldehyde-3-phosphate dehydrogenase is much smaller than the overall pool of 3PGA. This is also supported by the observation that species performing C_4_ photosynthesis are not characterised by high 3PGA/DHAP ratios (Fig. 4, see also Leegood & Furbank, 1984; Usuda, 1985; 1987; Leegood & von Caemmerer, 1988; 1989), even though the CCM in C_4_ photosynthesis requires additional energy (von Caemmerer & Furbank, 2003; Zhu *et al*., 2008). A third consistent feature with our proposal is that, apart from the high RuBP and 3PGA, the relative levels and relationships between metabolites in much of the rest of the CBC were in the range found in C_3_ and C_4_ plant species. Further evidence that high RuBP and 3PGA may serve to drive movement to and from carboxysome-located Rubisco is provided by the observation that RuBP and 3PGA levels were lower when the carboxysome-deficient Δ*ccmKLMN* mutant was compared with wild-type *Synechococcus* sp. PCC 7002 at 1% CO_2_ (Abernathy *et al*., 2019). This study also revealed that carboxysomes may play a wide-ranging role in the spatial micro-organisation of cyanobacterial metabolism. Küken *et al*. (2018) modelled fluxes between the stroma and pyrenoid matrix in *C. reinhardtii* and concluded that RuBP and 3PGA moved by diffusion in the absence of large concentration gradients of either metabolite. However, this conclusion was based on metabolite levels measured under very low irradiance and in high CO_2_. These conditions will have increased flux over the small fraction of Rubisco in the stroma, and decreased flux over Rubisco inside the pyrenoids and the need for metabolite gradients between the stroma and pyrenoid.

Some earlier more restricted studies also pointed to the presence of large pools of RuBP and 3PGA in cyanobacteria and algae at ambient CO_2_. Similar high levels for 3PGA (1300-3000 nmol g DW^-1^) and RuBP (200-300 nmol g DW^-1^) have been previously reported in *Synechocystis* (Takahashi *et al*., 2008; Marcus *et al*., 2011; Hasunuma *et al*., 2016; Hidese *et al*., 2021). Pioneering studies of metabolite profiles in *Chlorella pyrenoidosa* under ambient CO_2_ (Pedersen *et al*., 1966; Bassham and Krause, 1969) also reported relatively high RuBP. Interestingly, lower levels of RuBP have been reported under high CO_2_ in *C. pyrenoidosa* (Bassham and Kirk, 1960) and *C. reinhardtii* (Mettler *et al*., 2014), when pyrenoids partly disassemble. These observations are consistent with the high RuBP in ambient CO_2_ being partly due to location of Rubisco in the pyrenoid although it cannot be excluded that RuBP levels also decrease as a result of faster carboxylation in high CO_2_. It might be noted that the levels of 3PGA reported for *C. reinhardtii* in the present study are substantially higher than those previously reported (Mettler *et al*., 2014; Küken *et al*., 2018; Saint-Sorny *et al*., 2022). This is probably because these studies used reverse phase LC-MS/MS, when there can be considerable ion suppression of the 3PGA signal. The higher 3PGA levels reported in the present study were obtained using anion exchange LC-MS/MS where this complication is absent, and were confirmed by enzymatic assay.

In plants, the CBC is typically regulated such that Rubisco active sites are fully occupied by RuBP or, if this is not possible because photosynthesis is RuBP-limited, Rubisco is post-translationally inactivated (von Caemmerer & Farquhar, 1985; Woodrow & Berry, 1988; Sage *et al*., 1988; Parry *et al*., 2008; Sharkey, 2022). This is at least partly because other CBC metabolites would otherwise be bound by Rubisco catalytic sites, leading to sequestration of these metabolites and inactivation of Rubisco. In principle, the high RuBP content in cyanobacteria and algae might be explained if they have a high Rubisco content compared to C_3_ and C_4_ terrestrial plants, providing a higher concentration of binding sites in which RuBP is bound. However, this is very unlikely to be the case. Studies in cyanobacteria (Andrews & Abel, 1981; Read, 1994) and algae (Badger *et al*., 1998; Meyer & Griffiths, 2013; Heureux *et al*., 2017; Goudet *et al*., 2020) have shown that the high CO_2_ environment provided by their CCMs is accompanied by a change in Rubisco kinetics, with a relaxed selectivity for CO_2_ over O_2_ and a relatively high *k*_*cat*_, analogous to the situation in terrestrial C_4_ plants. For example, detailed studies of *C. reinhardtii* Rubisco revealed, compared to C_3_ and C_4_ plants, a higher K_m_CO_2_ (24-28 compared to 9-13 and 16-30 µM, respectively), similar or lower K_m_O_2_ (417-460 compared to 250-496 and 610 µM, respectively) and a higher V_max_ (119 compared to 63 µmol mg^-1^ protein in spinach) (Badger *et al*., 1998; Satagopan & Spreitzer, 2004; 2008; Sharwood *et al*., 2016; Rasineni *et al*., 2017). This higher *k*_*cat*_ allows allocation of nitrogen away from Rubisco and towards other proteins, analogous to the situation in terrestrial C_4_ species (see Introduction). At least in Chlamydomonas and in *Synechocystis*, Rubisco abundance (about 8.8% of total protein in Chlamydomonas, Hammel *et al*. (2020); about 2.15% of total protein in *Synechocystis*, Spaet *et al*. (2021)) are lower than in C_3_ plants and in the range found in C_4_ plants (see Sharwood *et al*., 2016).

Sequestration of Rubisco in a microcompartment may require adjustments in regulation of the remainder of the CBC (Fig. 5). In particular, compared to plants, cyanobacteria and algae not only operate with high RuBP levels (Fig. 3) but also with much lower pentose-P levels (Fig. 3) and, as a consequence, a much lower pentose-P/RuBP ratios (Fig. 4). Further, much of the RuBP will not be bound to Rubisco catalytic sites because it is probably located outside the carboxysome or pyrenoid (see above). This contrasts with the situation in plants, where much or all of the RuBP is bound to Rubisco catalytic sites (see Introduction). The implication is that poising of the various CBC reactions is modified in cyanobacteria and algae to favour PRK relative to other CBC enzymes and allow maintenance of a large free RuBP pool. Recently, Saint-Sorny *et al*. (2022) investigated the temporal kinetics of CBC metabolites after illumination of a Rubisco-deficient *C. reinhardtii* mutant in the absence or present of acetate. RuBP rose progressively over 5 min to levels of 7 and 2 nmol 10^−7^ cells in the absence and presence of acetate, respectively, whilst DHAP, S7P and pentose-P remained at very low levels (about 0.05, 0.01 and 0.12 without acetate, and 0.15, 0.075 and 0.2 nmol 10^−7^ cells with acetate). These observations point to the ability of *C. reinhardtii* PRK to produce RuBP even when there is a very large free pool of RuBP.

**Fig. 5.**
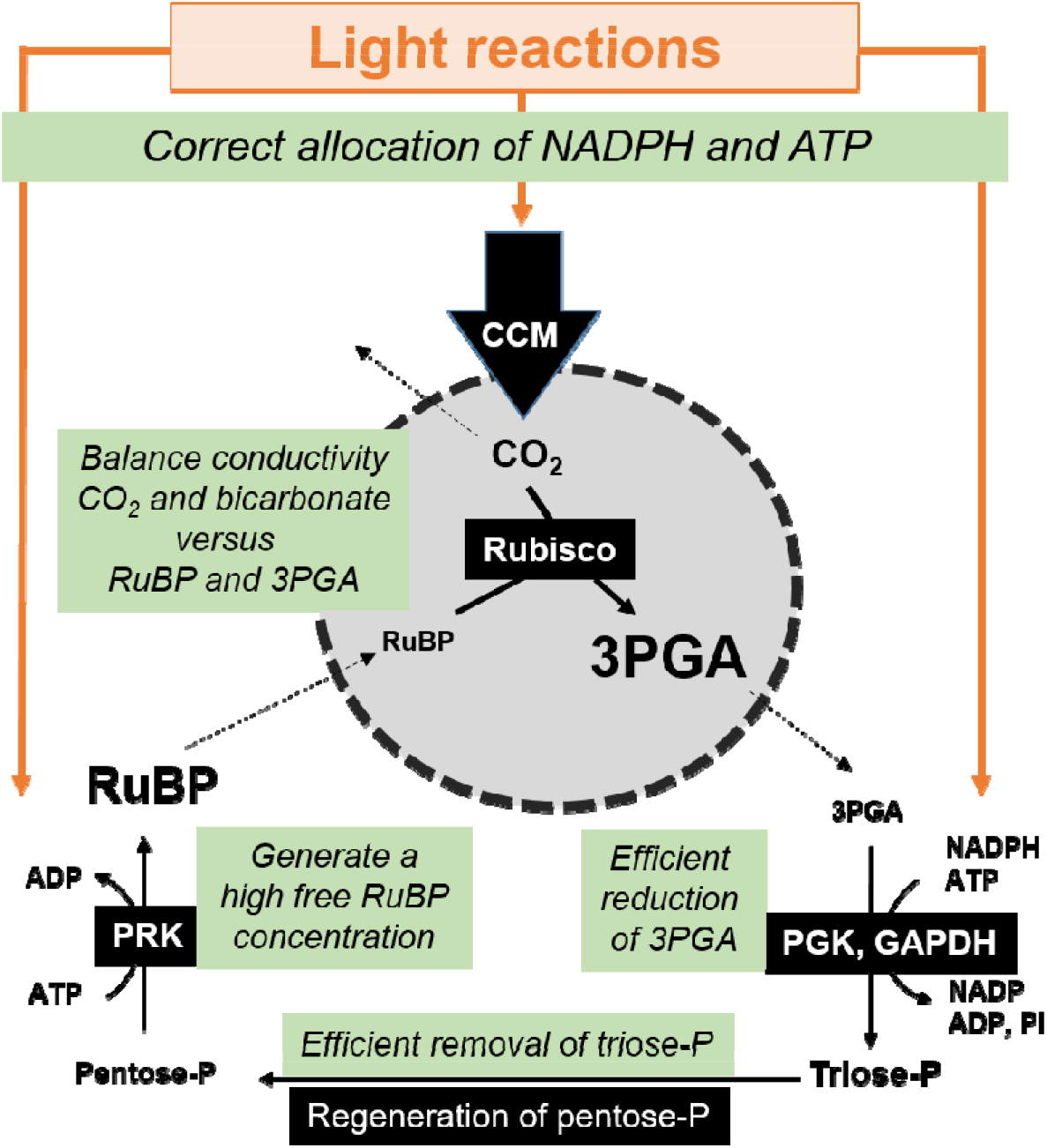
Scheme of features that may optimize CBC operation in conjunction with CCMs that require sequestration of Rubisco in a microcompartment. Operation of a cyanobacterial or algal CCM requires that the CBC generates high free RuBP to drive entry of RuBP into the microcompartment in which Rubisco is located. This will require regulation of PRK to generate these high RuBP concentrations, and of the remainder of the CBC to provide pentose-P for PRK. Movement of 3PGA out of the microcompartment will be aided by efficient reduction of 3PGA to triose-P. Coordination of the rate of CO_2_ concentration by the CCM will be important to maintain high enough CO_2_ concentrations in the CCM to suppress the oxygenation reaction of Rubisco (not shown) whilst not exceeding the rate of RuBP deliver and leading to excessive concentration and back-leakage of CO_2_. This will depend not only on the poising within the CBC but also on correct allocation of NADPH and ATP between the CCM, 3PGA reduction and the PRK reaction. Abbreviations: PGK = phosphoglycerate kinases; GAPDH = NADPH-dependent glyceraldehyde-3-phosphatedehydrogenase.

The strong propensity for RuBP synthesis in cyanobacteria and algae might be due to increased abundance or altered regulation of PRK protein. In plants, PRK is subject to strong feedback inhibition by downstream metabolites like RuBP, 3PGA and ADP (Gardemann *et al*., 1983; Stitt *et al*., 2010; 2021), with a fraction of the total PRK activity sufficing to catalyse CBC flux in steady state conditions (Paul *et al*., 2000; Stitt *et al*., 2010; Raines, 2011). It will be interesting to investigate if the regulatory characteristics of algal PRK differ from plant PRK, rendering it less susceptible to feedback inhibition by downstream metabolites, especially RuBP. The CBC metabolite profiles and metabolite ratios also indicate more resistance to flux at FBPase and SBPase in cyanobacteria and algae than in plants (Figs. 3-4). This might be a secondary consequence of increased capacity or deregulation of PRK. SBP levels and the SBP/S7P ratio were especially high in S*ynechocystis*, where hydrolysis of FBP and SBP is catalysed by a single bifunctional FBPase/SBPase (Tamoi *et al*., 1996; 1998; Yan & Xu, 2008). Overall, efficient reduction of 3PGA to triose-P and conversion of the latter to pentose-P and RuBP will favour movement of RuBP into and 3PGA out of the microcompartment (Fig. 5). Photosynthetic efficiency will be decreased if allocation of energy to 3PGA reduction and PRK is not correctly coordinated with allocation of energy to the CCM (Fig. 5). If the rate of concentration of CO_2_ lags behind consumption in the CBC this may result in higher rates of RuBP oxygenation, and if the rate of concentration of CO_2_ exceed consumption in the CBC there will be risk of increased back-leakage of CO_2_.

In conclusion, cyanobacteria and algae deploy CCMs that require sequestration of Rubisco in a microcompartment, with the consequence that Rubisco is physically separated from the remainder of the CBC enzymes. Using metabolite profiling, we have shown that, compared to terrestrial plants, cyanobacteria and algae exhibit a distinctive poising of the CBC, with high RuBP and high 3PGA levels. We propose that this is related to the need to drive diffusion of RuBP into and 3PGA out of the microcompartment that houses Rubisco. More studies are needed to identify physical features that may aid movement of these metabolites and to understand possible trade-offs with efficient concentration and retention of CO_2_ in these microcompartments. It can be envisaged that molecular changes in the carboxysome or pyrenoid surface that decrease conductivity for small molecules will decrease back-leakage of CO_2_ but might require even larger pools of RuBP and 3PGA to drive their movement into and out of the microcompartment. It will also be interesting to identify specific aspects of CBC regulation that favour its operation in conjunction with location of Rubisco in a microcompartment in cyanobacteria and algae. These will presumably have been required to co-evolve with the emergence of carboxysomes and pyrenoids, and that may have been relaxed or modified following loss of pyrenoids after colonisation of land masses and the subsequent evolution of terrestrial land plants. This highlights that operation of CCMs requires co-evolution of the CBC.

## Abbreviations

(2PG): 2-phosphoglycolate
(3PGA): 3-phosphoglycerate
(C): carbon
(CBC): Calvin-Benson cycle
(CCM): carbon-concentrating mechanism
(DHAP): dihydroxyacetone phosphate
(DW): dry weight
(FW): fresh weight
(F6P): fructose 6-phosphate
(FBPase): fructose-1,6-bisphosphatase
(FBP): fructose 1,6-bisphosphate
(GAP): glyceraldehyde 3-phosphate
(PEP): phospho*enol*pyruvate
(PRK): phosphoribulokinase
(pentose-P): pentose phosphates
(PC): principal component
(R5P): ribose 5-phosphate
(Ru5P): ribulose 5-phosphate
(RuBP): ribulose 1,5-bisphosphate
(Rubisco): ribulose 1,5-biphosphate carboxylase/oxygenase
(S7P): sedoheptulose 7-phosphate
(SBPase): sedoheptulose-1,7-bisphosphatase
(SBP): sedoheptulose 1,7-bisphosphate
(triose-P): triose phosphates
(Xu5P): xylulose 5-phosphate

## Acknowledgements

We acknowledge funding from the Max Planck Society (HT, RF, MS, SA), the HFSP (HT) and the German Research Foundation (DFG, HA 2002/23-1; SL, MH) as well as the University of Rostock (SL, MH).

## Author contributions

HT, SL, MS, MH and SA conceived and planned the experiments. HT and SL grew algae and cyanobacteria, respectively. HT and SL harvested and extracted the samples for downstream analyses. RF and SA processed the samples by LC-MS/MS. HT, RF and SA analysed the LC-MS/MS data. SA enzymatically assayed the samples. MS, MH and SA supervised the project. MS and SA wrote the manuscript. All authors approved the final version.

## Data availability

Data presented in this studies are reported in the Supporting Information Dataset S1.

## Supporting Information

**Table S1**. Comparison of 3PGA amounts determined by enzymatic assay and anion exchange LC-MS/MS.

**Table S2**. Summary of first 10 principal components (PC) for PCA analyses on cyanobacteria, algae and terrestrial plant species.

**Fig. S1**. Chromatograms of *Synechocystis* sp. PCC 6803 extract spiked with authentic GAP and DHAP standards.

**Fig. S2**. CBC metabolite profiles in terrestrial plant species.

**Fig. S3**. Principal components analysis of CBC metabolite profiles in terrestrial plant species.

**Fig. S4**. Metabolite ratios shown separately for cyanobacteria and algae (A) and for terrestrial plant species (B).

**Fig. S5**. Cross-species comparison of metabolite ratios of DHAP to other sugar phosphates.

**Method S1**. Additional information about cyanobacterial and algal cultures and harvest, sample and data processing.

**Dataset S1**. Metabolite levels and metabolite ratios in different species.

## References

Abernathy MH, Czajka JC, Allen DK, Hill NC, Camerond JC, Tang YT. 2019. Cyanobacterial carboxysome mutant analysis reveals the influence of enzyme compartmentalization on cellular metabolism and metabolic network rigidity. Metabolic Engineering 54, 222–231.

Andrews TJ, Abel KM. 1981. Kinetics and subunit interactions of ribulose bisphosphate carboxylase-oxygenase from the cyanobacterium, Synechococcus sp. Journal of Biological Chemistry 256, 8445–8451.

Arrivault S, Guenther M, Ivakov A, Feil R, Vosloh D, Van Dongen JT, Sulpice R, Stitt M. 2009. Use of reverse-phase liquid chromatography, linked to tandem mass spectrometry, to profile the Calvin cycle and other metabolic intermediates in Arabidopsis rosettes at different carbon dioxide concentrations. Plant Journal 59, 824–839.

Arrivault S, Guenther M, Fry SC, Fünfgeld MMFF, Veyel D, Mettler-Altmann T, Stitt M, Lunn JE. 2015. Synthesis and use of stable-isotope-labeled internal standards for quantification of phosphorylated metabolites by LC-MS/MS. Analytical Chemistry 87, 6896–6904.

Arrivault S, Alexandre Moraes T, Obata T, Medeiros DB, Fernie AR, Boulouis A, Ludwig M, Lunn JE, Borghi GL, Schlereth A et al. 2019. Metabolite profiles reveal interspecific variation in operation of the Calvin–Benson cycle in both C_4_ and C_3_ plants. Journal of Experimental Botany 70, 1843–1858.

Atkinson N, Velanis CN, Wunder T, Clarke DJ, Mueller-Cajar O, McCormick AJ. 2019. The pyrenoidal linker protein EPYC1 phase separates with hybrid Arabidopsis– Chlamydomonas Rubisco through interactions with the algal Rubisco small subunit. Journal of Experimental Botany 70, 5271–5285.

Atkinson N, Mao Y, Chan KX, McCormick AJ. 2020. Condensation of Rubisco into a proto-pyrenoid in higher plant chloroplasts. Nature Communications 11, 6303. https://doi.org/10.1038/s41467-020-20132-0

Badger MR, Price GD, Yu JW. 1991. Selection and analysis of mutants of the CO_2_-concentrating mechanism in cyanobacteria. Canadian Journal of Botany 69, 974–983.

Badger MR, Price GD. 1992. The CO_2_ concentrating mechanism in cyanobacteria and microalgae. Physiologia Plantarum 90, 529–536.

Badger MR, Andrews TJ, Whitney SM, Ludwig M, Yellowlees DC, Leggat W, Price GD. 1998. The diversity and coevolution of Rubisco, plastids, pyrenoids, and chloroplast-based CO_2_-concentrating mechanisms in algae. Canadian Journal of Botany 76, 1052–1071.

Badger MR, Price GD. 2003. CO_2_ concentrating mechanisms in cyanobacteria: molecular components, their diversity and evolution. Journal of Experimental Botany 54, 609–622.

Badger MR, Price GD, Long BM, Woodger FJ. 2006. The environmental plasticity and ecological genomics of the cyanobacterial CO_2_ concentrating mechanism. Journal of Experimental Botany 57, 249–265.

Bar-On YM, Milo R. 2019. The global mass and average rate of rubisco. Proceedings of the National Academy of Sciences of the United States of America 116, 4738–4743.

Barrett J, Girr P, Mackinder LCM. 2021. Pyrenoids: CO_2_-fixing phase separated liquid organelles. Biochimica Biophysica Acta - Molecular Cell Research 1868, https://doi.org/10.1016/j.bbamcr.2021.118949.

Bassham JA, Kirk M. 1960. Dynamics of the photosynthesis of carbon compounds I. Carboxylation reactions. Biochimica et Biophysica Acta 43, 447–464.

Bassham JA, Krause GH. 1969. Free energy changes and metabolic regulation in steady-state photosynthetic carbon reduction. Biochimica et Biophysica Acta 189, 207–221.

Bauwe H. 2018. Photorespiration – damage repair pathway of the Calvin Benson cycle. Annual Plant Reviews 50, 293–342.

Behrenfeld MJ, Randerson JT, McClain CR, Feldman GC, Los SO, Tucker CJ, Falkowski PG, Field CB, Frouin R, Esaias WE et al. 2001. Biospheric primary production during an ENSO transition. Science 291, 2594–2597.

Betti M, Bauwe H, Busch FA, Fernie AR, Keech O, Levey M, Ort DR, Parry MA, Sage R, Timm S et al. 2016. Manipulating photorespiration to increase plant productivity: Recent advances and perspectives for crop improvement. Journal of Experimental Botany 67, 2977–2988.

Blanco-Rivero A, Shutova T, Román MJ, Villarejo A, Martinez F. 2012. Phosphorylation controls the localization and activation of the lumenal carbonic anhydrase in Chlamydomonas reinhardtii. PLoS One 7, e49063. doi: 10.1371/journal.pone.0049063.

Borghi GL, Moraes TA, Günther M, Feil R, Mengin V, Lunn JE, Stitt M, Arrivault S. 2019. Relationship between irradiance and levels of Calvin-Benson cycle and other intermediates in the model eudicot Arabidopsis and the model monocot rice. Journal of Experimental Botany 70, 5809–5825.

Borghi GL, Arrivault S, Günther M, Barbosa Medeiros D, Dell’Aversana E, Fusco GM, Carillo P, Ludwig M, Fernie AR, Lunn JE, Stitt M. 2022. Metabolic profiles in C_3_, C_3_-C_4_ intermediate, C_4_-like, and C_4_ species in the genus Flaveria. Journal of Experimental Botany 73, 1581–1601.

Borkhsenious ON, Mason CB, J.V. Moroney JV. 1998. The intracellular localization of ribulose-1,5-bisphosphate carboxylase/oxygenase in Chlamydomonas reinhardtii. Plant Physiology 116, 1585–1591.

Bräutigam A, Schlüter U, Lundgren MR, Flachbart S, Ebenhöh O, Schönknecht G, Christin PA, Bleuler S, Droz JM, Osborne CP, Weber APM, U Gowik U. 2018. Biochemical mechanisms driving rapid fluxes in C_4_ photosynthesis. bioRxiv 387431; doi: https://doi.org/10.1101/387431

Brown RH. 1978. A difference in N use efficiency in C_3_ and C_4_ plants and its implications in adaptation and evolution. Crop Science 18, 93–98.

Burlacot A, Dao O, Auroy P, Cuiné S, Li-Beisson Y, Peltier G. 2022. Alternative photosynthesis pathways drive the algal CO_2_-concentrating mechanism. Nature 605, 366–371.

Busch FA. 2020. Photorespiration in the context of Rubisco biochemistry, CO_2_ diffusion and metabolism. The Plant Journal 101, 919–939.

Cai F, Sutter M, Cameron JC, Stanley DN, Kinney JN, Kerfeld CA. 2013. The structure of CcmP, a tandem bacterial microcompartment domain protein from the β-carboxysome, forms a subcompartment within a microcompartment. Journal of Biological Chemistry 288, 16055–16063.

Caspari OD, Meyer MT, Tolleter D, Wittkopp TM, Cunniffe NJ, Lawson T, Grossman AR, Griffiths G. 2017. Pyrenoid loss in Chlamydomonas reinhardtii causes limitations in CO_2_ supply, but not thylakoid operating efficiency. Journal of Experimental Botany 68, 3903–3913.

Cruz JA, Emery C, Wüst M, Kramer DM, Lange BM. 2008. Metabolite profiling of Calvin cycle intermediates by HPLC-MS using mixed-mode stationary phases. The Plant Journal 55, 1047–1060.

Dietz KJ, Heber U. 1984. Rate-limiting factors in leaf photosynthesis. I. Carbon fluxes in the calvin cycle. Biochimica Biophysica Acta - Bioenergetics 767, 432–443.

Doron L, Segal N and Shapira M. 2016. Transgene expression in microalgae—From tools to applications. Frontiers in Plant Science 7, 505. doi: 10.3389/fpls.2016.00505.

Ellis RJ. 1979. The most abundant protein in the world. Trends in Biochemical Sciences 4, 241–244.

Engel BD, Schaffer M, Kuhn Cuellar L, Villa E, Plitzko JM, Baumeister W. 2015. Native architecture of the Chlamydomonas chloroplast revealed by in situ cryo-electron tomography. Elife, 4:e04889. doi: 10.7554/eLife.04889.

Erb TJ, Zarzycki J. 2018. A short history of RuBisCO; the rise and fall(?) of Nature’s predominant CO_2_-fixing enzyme. Current Opinion in Biotechnology 49, 100–107.

Field CB, Behrenfeld MJ, Randerson JT, Falkowski P. 1998. Primary production of the biosphere: integrating terrestrial and oceanic components. Science 281, 237–240.

Flamholz AI, Prywes N, Moran U, Davidi D, Bar-On YM, Oltrogge LM, Alves R, Savage D, Milo R. 2019. Revisiting Trade-offs between Rubisco Kinetic Parameters. Biochemistry 58, 3365–3376.

Freeman Rosenzweig ES, Xu B, Kuhn Cuellar L, Martinez-Sanchez A, Schaffer M, Strauss M, Cartwright HN, Ronceray P, Plitzko JM, Förster F et al. 2017. The eukaryotic CO_2_-concentrating organelle is liquid-like and exhibits dynamic reorganization. Cell 171, 148–162.

Gardemann A, Stitt M, Heldt HW. 1983. Control of CO_2_ fixation. Regulation of spinach ribulose 5-phosphate kinase by stromal metabolite levels. Biochimica Biophysica Acta 722, 51–60.

Gerhardt R, Stitt M, Heldt HW. 1987. Subcellular metabolite levels in spinach leaves. Regulation of sucrose synthesis during diurnal alterations in photosynthesis. Plant Physiology 83, 399–407.

Giordano M, Beardall J, Raven JA. 2005. CO_2_ concentrating mechanisms in algae: mechanisms, environmental modulation, and evolution. Annual Review of Plant Biology 56, 99–131.

Goudet MMM, Orr DJ, Melkonian M, Müller KH, Meyer MT, Carma-Silva E, Griffiths H. 2020. Rubisco and carbon-concentrating mechanism co-evolution across chlorophyte and streptophyte green algae. New Phytologist 227, 810–823.

Griffiths H, Meyer MT, Rickaby REM. 2017. Overcoming adversity through diversity: aquatic carbon concentrating mechanisms. Journal of Experimental Botany 68, 3689–3695.

Hagemann M, Kern R, Maurino VG, Hanson DT, Weber APM, Sage RF, Bauwe H. 2016. Evolution of photorespiration from cyanobacteria to land plants, considering protein phylogenies and acquisition of carbon concentrating mechanisms. Journal of Experimental Botany 67, 2963–2976.

Hagemann M, Song S, Brouwer EM. 2021. Inorganic carbon assimilation in cyanobacteria: Mechanisms, regulation, and engineering. In Hudson P, Lee SY, Nielsen J (eds.) Cyanobacteria Biotechnology, Wiley-Blackwell Biotechnology Series, Chapter 1, 1–31, https://doi.org/10.1002/9783527824908.ch1

Hammel A, Sommer F, Zimmer D, Stitt M, Mühlhaus T, Schroda M. 2020. Overexpression of Sedoheptulose-1,7-Bisphosphatase enhances photosynthesis in Chlamydomonas reinhardtii and has no effect on the abundance of other Calvin-Benson cycle enzymes. Frontiers in Plant Science 11, 868. doi: 10.3389/fpls.2020.00868.

Hasunuma T, Harada K, Miyazawa SI, Kondo A, Fukusaki E, Miyake C. 2010. Metabolic turnover analysis by a combination of in vivo 13C-labelling from 13CO_2_ and metabolic profiling with CE-MS/MS reveals rate-limiting steps of the C_3_ photosynthetic pathway in Nicotiana tabacum leaves. Journal of Experimental Botany 61, 1041–1051.

Hasunuma T, Matsuda M, Kondo A. 2016. Improved sugar-free succinate production by Synechocystis sp. PCC 6803 following identification of the limiting steps in glycogen catabolism. Metabolic Engineering Communications 3, 130–141.

Hatch MD. 2002. C_4_ photosynthesis: discovery and resolution. Photosynthesis Research 73, 251–256.

Heldt HW, Piechulla B, Heldt F. 2005. Plant biochemistry. Cambridge, MA: Academic Press.

Heureux AMC, Young JN, Whitney SM, Eason-Hubbard MR, Lee RBY, Sharwood RE, Rickaby REM. 2017. The role of Rubisco kinetics and pyrenoid morphology in shaping the CCM of haptophyte microalgae. Journal of Experimental Botany 68, 3959–3969.

Hidese R, Matsuda M, Osanai T, Hasunuma T, Kondo A. 2020. Malic enzyme facilitates d-Lactate production through increased pyruvate supply during anoxic dark fermentation in Synechocystis sp. PCC 6803. ACS Synthetic Biology 9, 260–268.

Hohmann-Marriott MF, Blankenship RE. 2011. Evolution of photosynthesis. Annual Review of Plant Biology 62, 515–548.

Ikeuchi M, Tabata S. 2001. Synechocystis sp. PCC 6803 — a useful tool in the study of the genetics of cyanobacteria. Photosynthesis Research 70, 73–83.

Iniguez C, Capó-Bauçà S, Niinemets Ü, Stoll H, Aguiló-Nicolau P, Galmés J. 2020. Evolutionary trends in RuBisCO kinetics and their co-evolution with CO_2_ concentrating mechanisms. The Plant Journal 101, 897–918.

Itakura AK, Chan KX, Atkinson N, Pallesen L, Wang L, Reeves G, Patena W, Caspari O, Roth R, Goodenough U, McCormick AJ, Griffiths H, Jonikas MC. 2019. A Rubisco-binding protein is required for normal pyrenoid number and starch sheath morphology in Chlamydomonas reinhardtii. Proceedings of the National Academy of Sciences of the United States of America 116, 18445–18454.

Jordan DB, Ogren WL. 1981. Species variation in the specificity of ribulose biphosphate carboxylase/oxygenase. Nature 291, 513–515.

Kaplan A, Zenvirth D, Marcus Y, Omata T, Ogawa T. 1987. Energization and activation of inorganic carbon uptake by light in cyanobacteria. Plant Physiology 184, 210–213.

Kaplan A, Reinhold L. 1999. CO_2_ concentrating mechanisms in photosynthesis microorganisms. Annual Review of Plant Physiology and Plant Molecular Biology 50, 539–570.

Karlsson J, Clarke AK, Chen ZY, Hugghins SY, Park YI, Husic HD, Moroney JV, Samuelsson G. 1998. A novel alpha-type carbonic anhydrase associated with the thylakoid membrane in Chlamydomonas reinhardtii is required for growth at ambient CO_2_. The EMBO Journal 17, 1208–1216.

Kerfeld CA, Melnicki MR. 2016. Assembly, function and evolution of cyanobacterial carboxysomes. Current Opinion in Plant Biology 31, 66–75.

Kerfeld CA, Aussignargues C, Zarzycki J, Cai F, Sutter M. 2018. Bacterial microcompartments. Nature Reviews Microbiology 16, 277–290.

Küken A, Sommer F, Yaneva-Roder L, Höhne M, MacKInder L, Geimer S, Jonikas M, Schroda M, Stitt M, Nikoloski Z, Mettler-Altmann T. 2018. Effects of microcompartmentation on the distribution of fluxes and metabolic pools in the chloroplast of the green alga Chlamydomonas reinhardtii. eLIFE 2018;7:e37960. DOI: https://doi.org/10.7554/eLife.37960.

Kupriyanova EV, Sinetova MA, Cho SM, Park Y-I, Los DA, Pronina NA. 2013. CO_2_-concentrating mechanism in cyanobacterial photosynthesis: organization, physiological role, and evolutionary origin. Photosynthesis Research 117, 133–146.

Larsson AM, Hasse D, Valegård K, Andersson I. 2017. Crystal structures of β-carboxysome shell protein CcmP: ligand binding correlates with the closed or open central pore. Journal of Experimental Botany 68, 3857–3867.

Lechno-Yossef S, Brandon A. Rohnke BA, Belza ACO, Melnicki MR, Montgomery BL, Kerfeld CA. 2020. Cyanobacterial carboxysomes contain an unique rubisco-activase-like protein. New Phytologist 225, 793–806.

Leegood RC, Furbank RT. 1984. Carbon metabolism and gas exchange in leaves of Zea mays L.: Changes in CO_2_ fixation, chlorophyll a fluorescence and metabolite levels during photosynthetic induction. Planta 162, 450–456.

Leegood RC. 1985. The intercellular compartmentation of metabolites in leaves of Zea mays L. Planta 164, 163–171.

Leegood RC, von Caemmerer S. 1988. The relationship between contents of photosynthetic metabolites and the rate of photosynthetic carbon assimilation in leaves of Amaranthus edulis L. Planta 174, 253–262.

Leegood RC, von Caemmerer S. 1989. Some relationships between contents of photosynthetic intermediates and the rate of photosynthetic carbon assimilation in leaves of Zea mays L. Planta 178, 258–266.

Liu LN. 2022. Advances in the bacterial organelles for CO_2_ fixation. Trends in Microbiology 30, 567–580.

Long SP, Marshall-Colon A, Zhu XG. 2015. Meeting the global food demand of the future by engineering crop photosynthesis and yield potential. Cell 161, 56–66.

Lorimer GH. 1981. The carboxylation and oxygenation of Ribulose 1,5-Bisphosphate: The primary events in photosynthesis and photorespiration. Annual Review of Plant Physiology 32, 349–382.

Lunn JE, Feil R, Hendriks JH, Gibon Y, Morcuende R, Osuna D, Scheible WR, Carillo P, Hajirezaei MR, Stitt M. 2006. Sugar-induced increases in trehalose 6-phosphate are correlated with redox activation of ADPglucose pyrophosphorylase and higher rates of starch synthesis in Arabidopsis thaliana. Biochemical Journal 397, 139–148.

Ma F, Jazmin LJ, Young JD, Allen DK. 2014. Isotopically nonstationary 13C flux analysis of changes in Arabidopsis thaliana leaf metabolism due to high light acclimation. Proceedings of the National Academy of Sciences of the United States of America 111, 16967–16972.

Mackinder LCM, Meyer MT, Mettler-Altmann T, Chen V, Mitchell MC, Caspari O, Freeman Rosenzweig ES, Pallesen L, Reeves G, Itakura A et al. 2016. A repeat protein links Rubisco to form the eukaryotic carbon concentrating organelle. Proceedings of the National Academy of Sciences 113, 5958–5963.

Mackinder LCM, Chen C, Leib R, Patena W, Blum SR, Rodman M, Ramundo S, Adams C, Jonikas MC. 2017. A spatial interactome reveals the protein organization of the algal CO_2_ concentrating mechanism. Cell 171, 133–147.

Mann GD. 1996. Chloroplast morphology, movements and inheritance in diatoms. Cytology, Genetics and Molecular Biology of Algae, eds Chaudhary BR, Agrawal SB (SPB Academic Publishing, Amsterdam), 249–274.

Marcus Y, Berry JA, Pierce J. 1992. Photosynthesis and photorespiration in a mutant of the cyanobacterium Synechocystis PCC 6803 lacking carboxysomes. Planta 187, 511–516.

Marcus Y, Altman-Gueta H, Wolff Y, Gurevitz M. 2011. Rubisco mutagenesis provides new insight into limitations on photosynthesis and growth in Synechocystis PCC6803. Journal of Experimental Botany 62, 4173–4182.

Melnicki MR, Sutter M, Kerfeld CA. 2021. Evolutionary relationships among shell proteins of carboxysomes and metabolosomes. Current Opinion in Microbiology 63, 1–9.

Merlo L, Geigenberger P, Hajirezaei M, Stitt M. 1993. Changes of carbohydrates, metabolites and enzyme activities in potato tubers during development, and within a single tuber along astolon-apexgradient. Plant Physiology 142, 392–402.

Merchant SS, Prochnik SE, Vallon O, Harris EH, Karpowicz SJ, Witman GB, Terry A, Salamov A, Fritz-Laylin LK, Maréchal-Drouard L et al. 2007. The Chlamydomonas genome reveals the evolution of key animal and plant functions. Science 318, 245–25.

Mettler T, Mühlhaus T, Hemme D, Schöttler MA, Rupprecht J, Idoine A, Veyel D, Pal SK, Yaneva-Roder L, Winck FV et al. 2014. Systems analysis of the response of photosynthesis, metabolism, and growth to an increase in irradiance in the photosynthetic model organism Chlamydomonas reinhardtii. The Plant Cell 26, 2310–2350.

Meyer MT, Genkov T, Skepper JN, Jouhet J, Mitchell MC, Spreitzer RJ, Griffiths H. 2012. Rubisco small-subunit α-helices control pyrenoid formation in Chlamydomonas. Proceedings of the National Academy of Sciences of the United States of America 109, 19474–19479.

Meyer M, Griffiths H. 2013. Origins and diversity of eukaryotic CO_2_-concentrating mechanisms: lessons for the future. Journal of Experimental Botany 64, 769–786.

Meyer MT, Whittaker C, Griffiths H. 2017. The algal pyrenoid: key unanswered questions, Journal of Experimental Botany 68, 3739–3749.

Meyer MT, Itakura AK, Patena W, Wang L, He S, Emrich-Mills T, Lau CS, Yates G, Mackinder LCM, and Jonikas MC. 2020. Assembly of the algal CO_2_-fixing organelle, the pyrenoid, is guided by a Rubisco-binding motif. Science Advances 6, eabd2408. doi: 10.1126/sciadv.abd2408.

Mitra M, Mason CB, Xiao Y, Ynalvez RA, Lato SM, Moroney JV. 2005. The carbonic anhydrase gene families of Chlamydomonas reinhardtii. Canadian Journal of Botany 83, 780–795.

Moroney V, Ynalvez RA. 2007. Proposed carbon dioxide concentrating mechanism in Chlamydomonas reinhardtii. Eukaryotic Cell 6, 1251–1259.

Mukherjee A, Lau CS, Walker CE, Rai A, Prejean CI, Yates G, Emrich-Mills T, Lemoine SG, Vinyard DJ, Mackinder LCM, Moroney JV. 2019. Thylakoid localized bestrophin-like proteins are essential for the CO_2_ concentrating mechanism of Chlamydomonas reinhardtii. Proceedings of the National Academy of Sciences of the United States of America 116, 16915–16920.

Ort DR, Merchant SS, Alric J, Barkan A, Blankenship RE, Bock R, Croce R, Hanson MR, Hibberd JM, Long SP et al. 2015. Redesigning photosynthesis to sustainably meet global food and bioenergy demand. Proceedings of the National Academy of Sciences of the United States of America 112, 8529–8536.

Osmond CB. 1981. Photorespiration and photoinhibition: some implications for the energetics of photosynthesis. Biochimica et Biophysica Acta 639, 77–98.

Parry MAJ, Keys AJ, Madgwick PJ, Carmo-Silva AE, Andralojc PJ. 2008. RuBisCO regulation: A role for inhibitors. Journal of Experimental Botany 59, 1569–1580.

Paul MJ, Driscoll SP, Andralojc PJ, Knight JS, Gray JC, Lawlor DW. 2000. Decrease of phosphoribulokinase activity by antisense RNA intransgenic tobacco: definition of the light environment under which phosphoribulokinase is not in large excess. Planta 211, 112–119.

Pearce FG. 2006. Catalytic by-product formation and ligand binding by ribulose bisphosphate carboxylases from different phylogenies. Biochemical Journal 399, 525–534.

Pedersen TA, Kirk M, Bassham JA. 1966. Light-Dark transients in levels of intermediate compounds during photosynthesis in air-adapted Chlorella. Physiologia Plantarum 19, 219–231.

Price GD, Pengelly JJ, Forster B, Du J, Whitney SM, von Caemmerer S, Badger MR, Howitt SM, Evans JR. 2013. The cyanobacterial CCM as a source of genes for improving photosynthetic CO_2_ fixation in crop species. Journal of Experimental Botany 64, 753–768.

Rae BD, Long BM, Badger MR, Price GD. 2013. Functions, compositions, and evolution of the two types of carboxysomes: polyhedral microcompartments that facilitate CO_2_ fixation in cyanobacteria and some proteobacteria. Microbiology and Molecular Biology Reviews 77, 357–379.

Raines CA. 2011. Increasing photosynthetic carbon assimilation in C_3_ plants to improve crop yield: current and future strategies. Plant Physiology 155, 36–42.

Rasineni GK, Loh PC, Lim BH. 2017. Characterization of Chlamydomonas Ribulose-1,5-bisphosphate carboxylase/oxygenase variants mutated at residues that are post-translationally modified. Biochimica et Biophysica Acta 1861, 79–85.

Rasmussen B, Fletcher IR, Brocks JJ, Kilburn MR. 2008. Reassessing the first appearance of eukaryotes and cyanobacteria. Nature 455, 1101–1104.

Raven JA, Beardall J, Sánchez-Baracaldo P. 2017. The possible evolution and future of CO_2_-concentrating mechanisms. Journal of Experimental Botany 68, 3701–3716.

Read BA. 1994. High substrate specificity factor ribulose bisphosphate carboxylase/oxygenase from eukaryotic marine algae and properties of recombinant cyanobacterial rubisco containing “algal” residue modifications. Archives of Biochemistry and Biophysics 312, 210–218.

Rippka R, Deruelles J, Waterbury JB, Herdman M, Stanier RY. 1979. Generic assignments, strain histories and properties of pure cultures of cyanobacteria. Journal of General Microbiology 111, 1–61.

Rochaix JD. 2002. Chlamydomonas, a model system for studying the assembly and dynamics of photosynthetic complexes, FEBS Letters 529, 34–38.

Rousseaux CS, Gregg WW. 2013. Interannual variation in phytoplankton primary production at a global scale. Remote Sensing 6, 1–19.

Ryu Y, Berry JA, Baldocchi DD. 2019. What is global photosynthesis? History, uncertainties and opportunities. Remote Sensing of the Environment 223, 95–114.

Sage RF, Sharkey TD, Seemann JR. 1988. The in vivo response of ribulose-1,5-bisphosphate carboxylase activation state and pool sizes of photosynthetic metabolites to elevated CO_2_ in Phaseolus vulgaris L. Planta 174, 407–416.

Sage RF, Sage TL, Kocacinar F. 2012. Photorespiration and the evolution of C4 photosynthesis. Annual Review of Plant Biology 63, 19–47.

Sage RF. 2017. A portrait of the C_4_ photosynthetic family on the 50th anniversary of its discovery: species number, evolutionary lineages, and Hall of Fame. Journal of Experimental Botany 68, 4039–4056.

Saint-Sorny M, Brzezowski P, Arrivault S, Alricand J, Johnson X. 2022. Interactions between carbon metabolism and photosynthetic electron transport in a Chlamydomonas reinhardtii mutant without CO_2_ fixation by RuBisCO. Frontiers in Plant science https://doi.org/10.3389/fpls.2022.876439

Salome PA, Merchant SS. 2019. A series of fortunate events: Introducing Chlamydomonas as a reference organism. The Plant Cell 31, 1682–1707.

Salvucci ME. 1989. Regulation of Rubisco activity in vivo. Plant Physiology 77, 164–171.

Satagopan S, Spreitzer RJ. 2004. Substitutions at the Asp-473 latch residue of chlamydomonas ribulosebisphosphate carboxylase/oxygenase cause decreases in carboxylation efficiency and CO_2_/O_2_ specificity. Journal of Biological Chemistry 14, 14240–14244.

Satagopan S, Spreitzer RJ. 2008. Plant-like substitutions in the large-subunit carboxy terminus of Chlamydomonas Rubisco increase CO_2_/O_2_ specificity. BMC Plant Biology 30, 8–85.

Savir Y, Noor E, Milo R, Tluty T. 2010. Cross-species analysis traces adaptation of rubisco towards optimality in a low-dimensional landscale. Proceedings of the National Academy of Sciences 197, 3478–3480.

Sharkey TD. 1988. Estimating the rate of photorespiration in leaves. Physiologia Plantarum 73, 147–152.

Sharkey TD. 2022. The discovery of Rubisco. Journal of Experimental Botany, in press.

Sharwood RE, Sonawane BV, Ghannoum O, Whitney SM. 2016. Improved analysis of C_4_ and C_3_ photosynthesis via refined in vitro assays of their carbon fixation biochemistry. Journal of Experimental Botany 67, 3137–3148.

Shih PM, Occhialini A, Cameron JC, Andralojc PJ, Parry MAJ, Kerfeld CA. 2016. Biochemical characterization of predicted Precambrian RuBisCO. Nature Communications 7, 10382. https://doi.org/10.1038/ncomms10382.

Sinetova MA, Kupriyanova EV, Markelova AG, Allakhverdiev SI, Pronina NA. 2012. Identification and functional role of the carbonic anhydrase Cah3 in thylakoid membranes of pyrenoid of Chlamydomonas reinhardtii. Biochimica et Biophysica Acta 1817, 1248–1255.

So AKC, John-McKay M, Espie GS. 2002. Characterization of a mutant lacking carboxysomal carbonic anhydrase from the cyanobacterium Synechocystis PCC6803. Planta 214, 456–467.

Spalding MH. 2008. Microalgal carbon-dioxide-concentrating mechanisms: Chlamydomonas inorganic carbon transporters. Journal of Experimental Botany 59, 1463–1473.

Stitt M, Wirtz W, Heldt HW. 1980. Metabolite levels in the chloroplast and extrachloroplast compartments of spinach protoplasts. Biochimica et Biophysica Acta 593, 85–102.

Stitt M, Heldt HW. 1985. Generation and maintenance of concentration gradients between the mesophyll and bundle sheath in maize leaves. Biochimica et Biophysica Acta 808, 400– 414.

Stitt M, Lunn J, Usadel B. 2010. Arabidopsis and primary photosynthetic metabolism— more than the icing on the cake. The Plant Journal 61, 1067–1091.

Stitt M, Borghi GL, Arrivault S. 2021. Targeted metabolite profiling as a top-down approach to uncover interspecies diversity and identify key conserved operational features in the Calvin-Benson cycle. Journal of Experimental Botany 72, 5961–5986.

Sun Y, Wollman AJM, Huang F, Leake MC, Liu LN. 2019. Single-organelle quantification reveals stoichiometric and structural variability of carboxysomes dependent on the environment. Plant Cell 31, 1648–1664.

Takahashi H, Uchimiya H, Hihara Y. 2008. Difference in metabolite levels between photoautotrophic and photomixotrophic cultures of Synechocystis sp. PCC 6803 examined by capillary electrophoresis electrospray ionization mass spectrometry. Journal of Experimental Botany 59, 3009–3018.

Tamoi M, Ishikawa T, Takeda T, Shigeoka S. 1996. Molecular characterization and resistance to hydrogen peroxide of two fructose-1,6-bisphosphatases from Synechococcus PCC 7942. Archives of Biochemistry and Biophysics 334, 27–36.

Tamoi M, Murakami A, Takeda T, Shigeoka S. 1998. Acquisition of a new type fructose-1,6-bisphosphatase with resistance to hydrogen peroxide in cyanobacteria: molecular characterization of the enzyme from Synechocystis PCC 6803. Biochimica et Biophysica Acta 1383, 232–244.

Tcherkez GGB, Farquhar GD, Andrews TJ. 2006. Despite slow catalysis and confused substrate specificity, all ribulose bisphosphate carboxylases may be nearly perfectly optimized. Proceedings of the National Academy of Sciences of the United States of America 103, 7246–7251.

Thierstein HR, Young JR. 2004. Coccolithophores: From molecular processes to global impact (Springer, Heidelberg, Germany). DOI:10.1007/978-3-662-06278-4.

Toyokawa C, Yamano T, Fukuzawa H. 2020. Pyrenoid Starch sheath is required for LCIB localization and the CO_2_-concentrating mechanism in green algae. Plant Physiology 182, 1883–1893.

Treves H, Raanan H, Finkel OM, Berkowicz SM, Keren N, Shotland Y, Kaplan A. 2013. A newly isolated Chlorella sp. from desert sand crusts exhibits a unique resistance to excess light intensity. FEMS Microbiology Ecology 86, 373–80.

Treves H, Murik O, Kedem I, Eisenstadt D, Meir S, Rogachev I, Szymanski J, Keren N, Orf I, Tiburcio AF et al. 2017. Metabolic flexibility underpins growth capabilities of the fastest growing alga. Current Biology 27, 2559–2567.

Treves H, Siemiatkowska B, Luzarowska U, Murik O, Fernandez-Pozo N, Moraes TA, Erban A, Armbruster U, Brotman Y, Kopka J et al. 2020. Multi-omics reveals mechanisms of total resistance to extreme illumination of a desert alga. Nature Plants 6, 1031–1043.

Treves H, Arrivault S, Ishihara H, Hoppe I, Erban A, Kopka J, Hoehne M, Moraes TA, Szymanski J, Nikoloski Z, Stitt M. 2022. Carbon flux through photosynthesis and central carbon metabolism show distinct patterns between algae, C_3_ and C_4_ plants. Nature Plants 8, 78–91.

Usuda H. 1985. Changes in levels of intermediates of the C_4_ cycle and reductive Pentose Phosphate Pathway during induction of photosynthesis in maize leaves. Plant Physiology 78, 859–864.

Usuda H. 1987. Changes in levels of intermediates of the C_4_ cycle and reductive Pentose Phosphate Pathway under various light intensities in maize leaves. Plant Physiology 84, 549–554.

von Caemmerer S, Farquhar GD. 1985. Kinetics and activation of Rubisco and some preliminary modelling of the RuP2 pool sizes. In J Viil, G Grishina, A Laisk, eds, Kinetics of photosynthetic carbon metabolism in C3-plants. Estonian Academy of Sciences, Tallin, pp 46-58.

von Caemmerer S, Furbank R. 2003. The C_4_ pathway: an efficient CO_2_ pump. Photosynthesis Research 77, 191–207.

Wang Y, Spalding MH. 2014. Acclimation to very low CO_2_: Contribution of limiting CO_2_ inducible proteins, LCIB and LCIA, to inorganic carbon uptake in Chlamydomonas reinhardtii. Plant Physiology 166, 2040–2050.

Wang Y, Stessman DJ, Spalding MH. 2015. The CO_2_ concentrating mechanism and photosynthetic carbon assimilation in limiting CO_2_: how Chlamydomonas works against the gradient. Plant Journal 82, 429–448.

Woodrow IE, Berry JA. 1988. Enzymatic regulation of photosynthetic CO_2_ fixation in C_3_ plants. Annual Review of Plant Physiology and Plant Molecular Biology 39, 533–594.

Xu M, Bernát G, Singh A, Mi H, Rögner M, Pakrasi HB, Ogawa T. 2008. Properties of mutants of Synechocystis sp. strain PCC 6803 lacking inorganic carbon sequestration systems. Plant Cell Physiology 49, 1672–1677.

Yan C, Xu X. 2008. Bifunctional enzyme FBPase/SBPase is essential for photoautotrophic growth in cyanobacterium Synechocystis sp. PCC 6803. Progress in Natural Science 18, 149–153.

Yamano T, Sato E, Iguchi H, Fukuda Y, Fukuzawa H. 2015. Characterization of cooperative bicarbonate uptake into chloroplast stroma in the green alga Chlamydomonas reinhardtii. Proceedings of the National Academy of Sciences of the United States of America 112, 7315–7320.

Yeoh HH, Badger MR, Watson L. 1980. Variations in KmCO_2_ of Ribulose-1,5-bisphosphate carboxylase among grasses. Plant Physiology 66, 1110–1112.

Young JN, Heureux AMC, Sharwood RE, Rickaby REM, Morel FMM, Whitney SM. 2016. Large variation in the Rubisco kinetics of diatoms reveals diversity among their carbon-concentrating mechanisms. Journal of Experimental Botany 67, 3445–3456.

Zachos JC, Dickens GR, Zeebe RE. 2008. An early Cenozoic perspective on greenhouse warming and carbon-cycle dynamics. Nature 451, 279–283.

Zhan Y, Marchand CH, Maes A, Mauries A, Sun Y, Dhaliwal JS, Uniacke J, Arragain S, Jiang H, Gold ND et al. 2018. Pyrenoid functions revealed by proteomics in Chlamydomonas reinhardtii. PLoS ONE 13, e0185039. doi: 10.1371/journal.pone.0185039

Zhu XG, Long SP, Ort DR. 2008. What is the maximum efficiency with which photosynthesis can convert solar energy into biomass? Current Opinion in Biotechnology 19, 153–159.

