## Supplementary material for "Operation of Carbon-Concentrating Mechanisms in Cyanobacteria and Algae requires altered poising of the Calvin-Benson cycle"

The following Supporting Information is available for this article:

**Table S1.** Comparison of 3PGA amounts determined by enzymatic assay and anion exchange LC-MS/MS.

**Table S2.** Summary of first 10 principal components (PC) for PCA analyses on cyanobacteria, algae and terrestrial plant species.

**Fig. S1.** Chromatograms of *Synechocystis* sp. PCC 6803 extract spiked with authentic GAP and DHAP standards.

**Fig. S2.** CBC metabolite profiles in terrestrial plant species.

**Fig. S3.** Principal components analysis of CBC metabolite profiles in terrestrial plant species.

**Fig. S4.** Metabolite ratios shown separately for cyanobacteria and algae (A) and for terrestrial plant species (B).

**Fig. S5.** Cross-species comparison of metabolite ratios of DHAP to other sugar phosphates.

**Method S1.** Additional information about cyanobacterial and algal cultures and harvest, sample and data processing.

**Dataset S1.** Metabolite levels and metabolite ratios in different species (additional file).

**Table S1. Comparison of 3PGA amounts determined by enzymatic assay and anion exchange LC-MS/MS.** Measurements were done with LC-MS/MS extracts and values are expressed as nmol g DW^-1^. Additional samples (indicated with an asterisk) were obtained for 3PGA quantification. The absence of an asterisk indicates that the sample is used in this study. The ratios of 3PGA determined by enzymatic assay and 3PGA determined by LC-MS/MS are also presented.

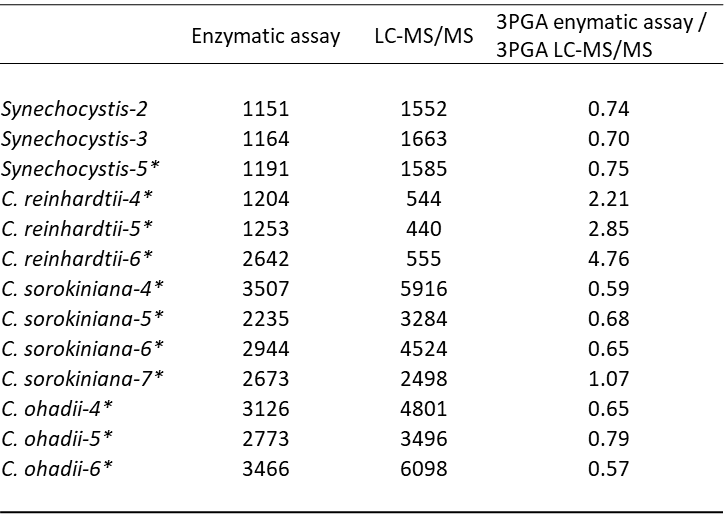

**Table S2. Summary of first 10 principal components (PC) for PCA analyses on cyanobacteria, algae and terrestrial plant species.** This Table is supplemental to Fig. 2.
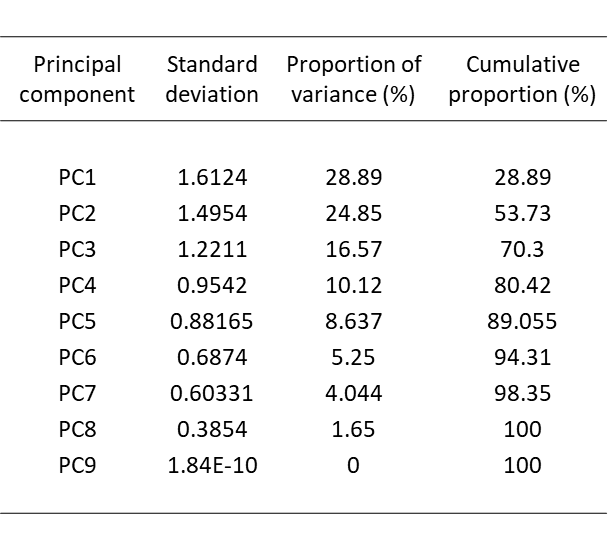

**Fig. S1. Chromatograms of *Synechocystis* sp. PCC 6803 extract spiked with authentic GAP and DHAP standards. (A**) Extracts not spiked, **(B)** extract spiked with GAP, **(C)** extracts spiked with DHAP, **(D)** GAP standard, **(E)** DHAP standards and **(F)** GAP and DHAP standards. Twenty pmol of standards were used. Retention times are indicated (in min).

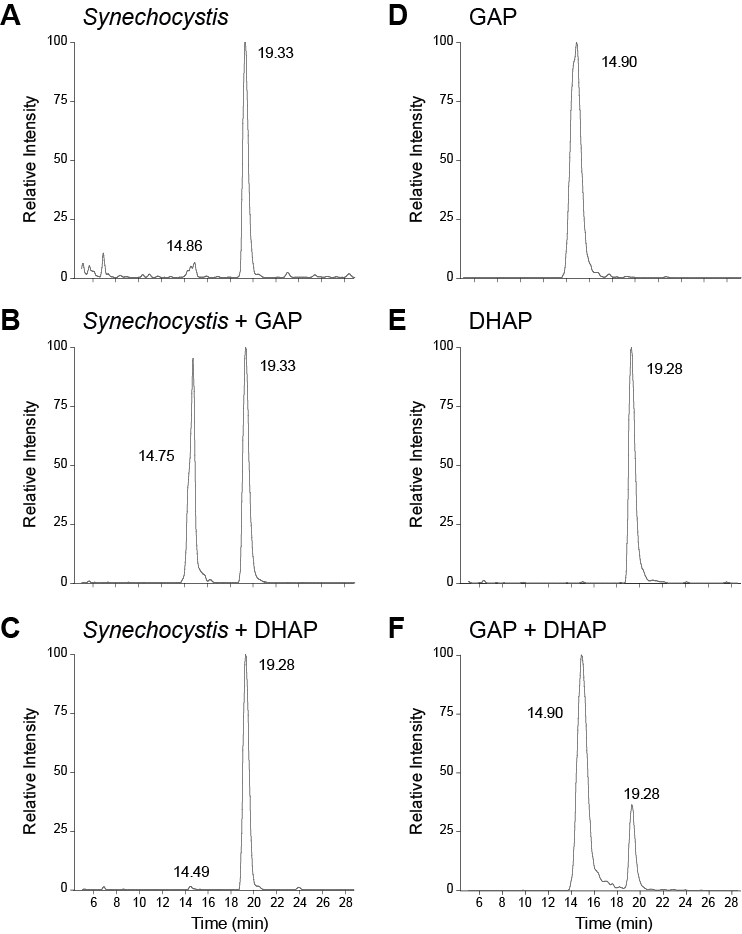

**Fig. S2. CBC metabolite profiles in terrestrial plant species.** The data are replotted from Arrivault *et al*. (2019) and Borghi *et al.* (2021). Species are ordered from left to right, C_3_, C_3_-C_4_ intermediate, C_4_-like and C_4_ terrestrial plant species. The results are shown as mean (nmol g FW^-1^) ± SD (*n*=3 to 9). Data are presented in Dataset S1.

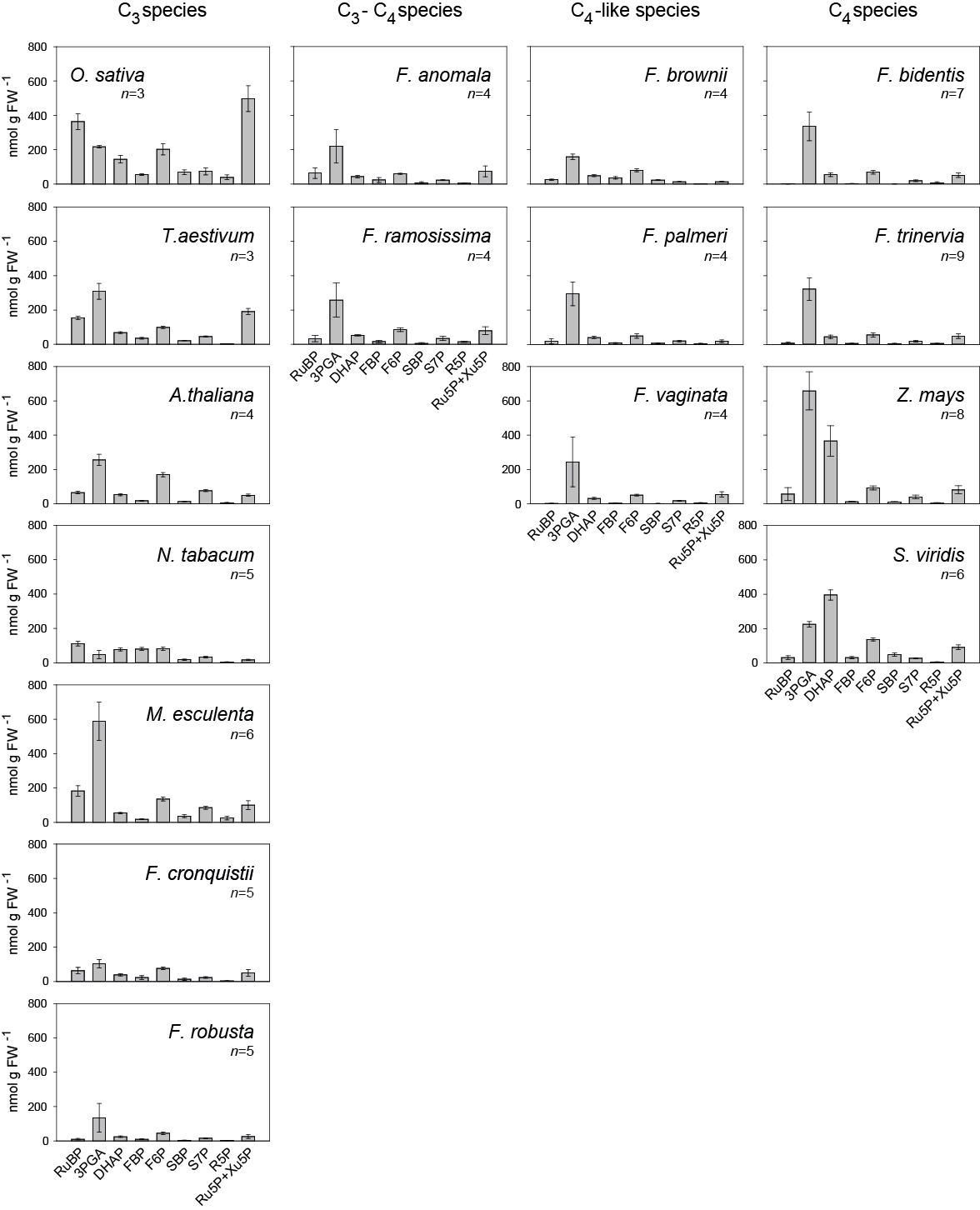

**Fig. S3. Principal components analysis of CBC metabolite profiles in terrestrial plant species.** The data are replotted from Arrivault *et al.* (2019) and Borghi *et al*. (2021), excluding 2PG. Terrestrial C_3_, C_4_, C_3_-C_4_ intermediate and C_4_-like plants are denoted by colour (black, grey, pale green and green, respectively; see insert legend). Abbreviations: Species abbreviations, full name and photosynthesis mode are, alphabetically: At, *Arabidopsis thaliana* (C_3_); Fan, *Flaveria anomala* (C_3_-C_4_); Fbi, *Flaveria bidentis* (C_4_); Fbr, *Flaveria brownii* (C_4_-like); Fcr, *Flaveria cronquistii* (C_3_); Fpa, *Flaveria palmeri* (C_4_-like); Fra, *Flaveria ramosissima* (C_3_-C_4_); Fro, *Flaveria robusta* (C_3_); Ftr, *Flaveria trinervia* (C_4_); Fva, *Flaveria vaginata* (C_4_-like); Me, *Manihot esculenta* (C_3_); Nt, *Nicotiana tabacum* (C_3_); Os, *Oryza sativa* (C_3_); Sv, *Setaria viridis* (C_4_); Ta, *Triticum aestivum* (C_3_); Zm, *Zea mays* (C_4_). Data are presented in Dataset S1, including terrestrial plant data taken from Arrivault *et al.* (2019) and Borghi *et al*. (2021).

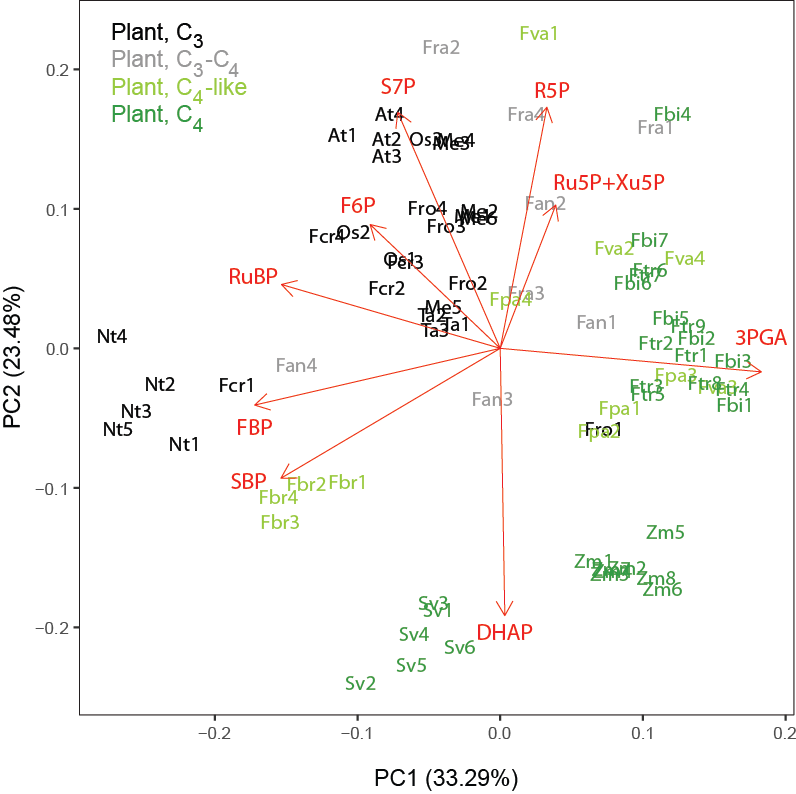

**Fig. S4. Metabolite ratios shown separately for cyanobacteria and algae (A) and for terrestrial plant species (B)**. This Figure is supplemental to Fig. 4. Ratios were determined using metabolite amounts. The data are displayed as box plots (*n*=3 to 9). Species are ordered from left to right, cyanobacteria (Cy, purple) and pyrenoidal eukaryotic algae (blue) in **(A)**, and C_3_ (black), C_3_-C_4_ intermediate (grey), C_4_-like (light green) and C_4_ terrestrial plant species (green) in **(B)**. For species abbreviations see legend of Fig. 2.

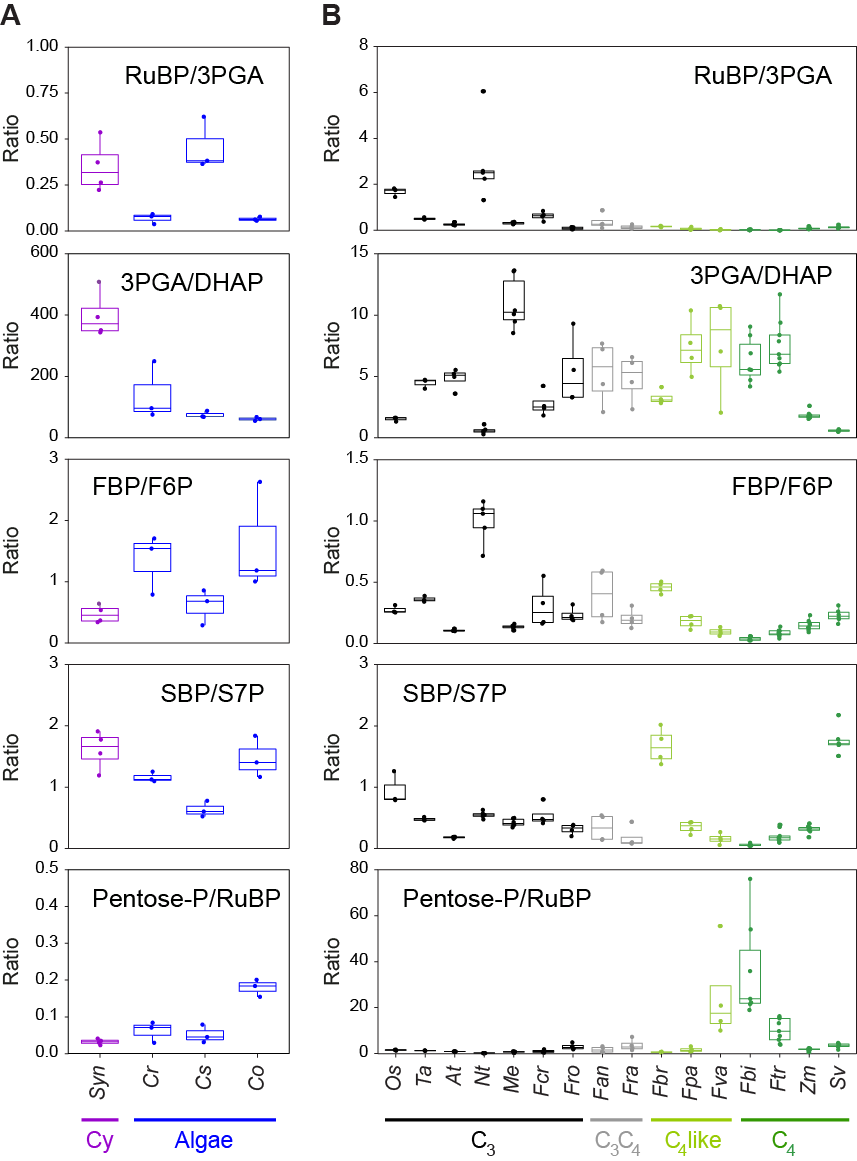

**Fig. S5. Cross-species comparison of metabolite ratios of DHAP to other sugar phosphates.** This Figure is supplemental to Fig. 4. Ratios were determined using metabolite amounts. The data are displayed as box plots (*n*=3 to 9). Species are ordered from left to right, cyanobacteria (Cy, purple), pyrenoidal eukaryotic algae (blue), C_3_ (black), C_3_-C_4_ intermediate (grey), C_4_-like (light green) and C_4_ terrestrial plant species (green). In case of need for a better visualization, inserts with cyanobacteria and algae species are included. For species abbreviations see legend of Fig. 2. Ratios are presented in Dataset S1, including terrestrial plant data taken from Arrivault *et al.* (2019) and Borghi *et al*. (2021).

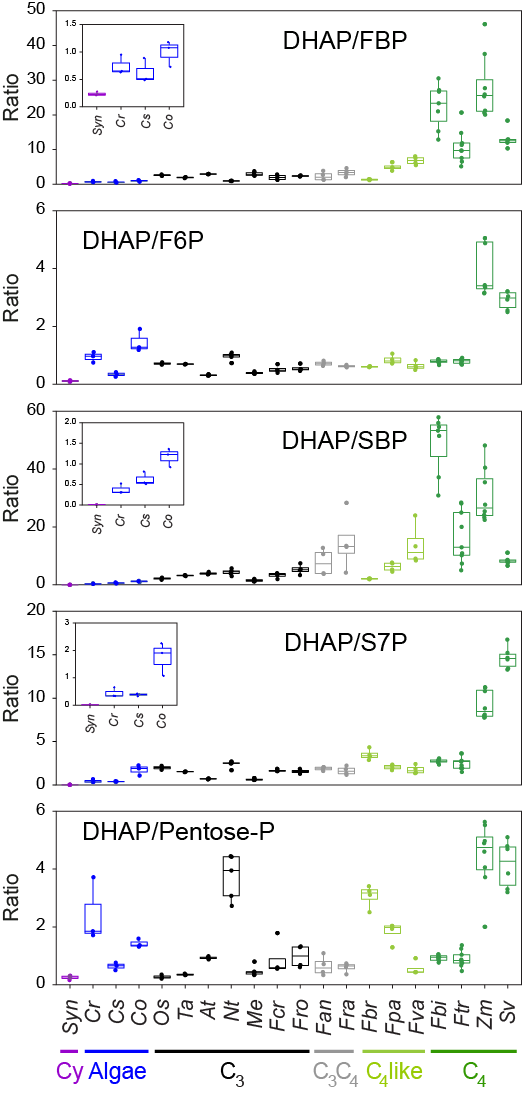

**Method S1. Additional information about cyanobacterial and algal cultures and harvest, sample and data processing.**

***Material***

N_2_, O_2_ and CO_2_ were obtained from Air Liquide (Germany; www.industrie.airliquide.de), and chemicals from Sigma-Aldrich (Darmstadt, Germany; www.sigmaaldrich.com), Roche Applied Science (Mannheim, Germany; lifescience.roche.com) or Merck (www.merckmillipore.com).

***Cyanobacterial and algal cultures and harvests***

Cultures of *Synechocystis* sp*.* PCC 6803 were grown in glass tubes supplied with air bubbling in buffered BG-11 medium (Rippka *et al*., 1979; TES pH 8.0) at 30°C and 100 µmol photons m^-2^ s^-1^. Cells were pre-cultivated at ambient air for one week, then harvested by centrifugation (5000 *g*, 7 min, 20°C) and subsequently resuspended in fresh medium to OD_750nm_ = 1.0. These cells were grown for another 24 h. Before the sampling, the cultures were then similarly adjusted to OD_750nm_ = 1.0 and allowed to reacclimate to the growth conditions for another three hours. Following the measurement of OD_750nm_, samples of 8 mL were harvested with glass pipettes in at least three technical triplicates per culture and quenched directly in 16 mL 70% methanol solution cooled to -70^o^C taking care that this occurred under growth irradiance. Quenched cells were then harvested by centrifugation (10000 *g*, 7 min, 4°C), supernatants discarded and the pellets frozen in liquid nitrogen and stored at -80°C until extraction.

Cultures of *C. ohadii*, *C. sorokiniana* and *C. reinhardtii* were grown in flat glass bioreactor vessels (FMT-150, PSI, Drasov, Czech Republic). The medium in the bioreactors was HP without acetate (Mettler *et al.*, 2014). Experiments were initiated at low [cell density](https://www.sciencedirect.com/topics/biochemistry-genetics-and-molecular-biology/cell-density) corresponding to OD_735nm_ = 0.02 and the culture temperature was stabilized at 35.0 ± 0.3°C for *C. ohadii* and *C. sorokiniana* or 25.0 ± 0.3°C for *C. reinhardtii*. Irradiance level was 100 μmol photons m^−2^ s^−1^, and air for bubbling was supplied using an air-pump at ∼1 L min^-1^. Bioreactors were autoclaved with medium and electrodes, and axenic inoculum was added on a sterile bench, together with 0.22 µm filters on all air inlets/outlets of the system. The cultures were axenic, as validated with light microscopy (Eclipse E200, Nikon, Melville, NY, USA) and LB plating/incubation. Independent biological replicates were collected from three separate bioreactor runs for each alga or condition. Cultures and fresh media were pushed through a tailor-made transparent mixer into 70% methanol solution cooled to -70^o^C as described in Treves *et al*. (2022), taking care that quenching occurred under growth irradiance. Quenched samples were centrifuged (3200 *g*, 3 min, -9^o^C), supernatants discarded and the pellets frozen in liquid nitrogen and stored at -80°C until extraction.

***Metabolite extraction and analysis***

Metabolites were extracted from cyanobacteria and algae as in Mettler *et al.* (2014) with some modifications of the first steps. Cyanobacterial frozen pellets were resuspended in -20°C cold 47% methanol and glass beads were not used for the three cycles of freeze-thaw. Algae frozen pellets were resuspended in -20°C cold methanol/chloroform (5:1, v/v) followed by four cycles of freeze-thaw. Metabolites were quantified by liquid chromatography linked to tandem mass spectrometry (LC-MS/MS) using either reverse phase LC (Arrivault *et al.*, 2009; all CBC metabolites except 3PGA) or anion exchange LC (Lunn *et al.,* 2006) for 3PGA (in the reverse phase platform 3PGA can be subject to peak spreading and ion suppression) . Samples were spiked with stable-isotope-labelled standards to correct for ion suppression and other matrix effects (Arrivault *et al.*, 2015). 3PGA was also quantified enzymatically in LC-MS/MS extracts (Merlo *et al.,* 1993).

**Statistical Analysis**

Histograms were prepared in Sigma Plot. Box plots and principal component analysis were performed in R Studio using ggpubr (ggplot2; <https://cran.r-project.org/web/packages/ggpubr/index.html>) and ggfortify (<https://cran.r-project.org/web/packages/ggfortify/index.html>) packages, respectively. Metabolite data (which were initially expressed on a dry weight basis for cyanobacteria and algae and a fresh weight basis for the published data on terrestrial plant species) were normalised. For each sample, the amount of C in a given metabolite was divided by the total amount of C in all CBC intermediates (as described in Arrivault *et al.*, 2019; see also Dataset S1).

Note. References listed in Method S1 are also listed in the main text,
